# Deaminase-based RNA recording enables high throughput mutational profiling of protein-RNA interactions

**DOI:** 10.1101/2025.04.11.648485

**Authors:** Rachael A. Bakker, Oliver B. Nicholson, Heungwon Park, Yu-Lan Xiao, Weixin Tang, Arvind Rasi Subramaniam, Christopher P. Lapointe

## Abstract

Protein-RNA interactions regulate RNA metabolism and are frequently dysregulated in disease. Scalable methods are needed to connect RNA and protein sequence requirements across biochemical and cellular settings. Here, we use the engineered TadA8.20 deaminase to record protein-RNA recognition as nucleotide edits read by DNA sequencing. Using the λN-boxB complex, we establish a common recording readout for three complementary comparisons. Pooled mutational scanning on both the RNA and protein sides recovers key boxB nucleotides and λN residues involved in recognition. Comparison of boxB libraries in vitro and in human cells reveals conserved sequence requirements for protein-RNA recognition. Direct fusion and split recruitment of deaminase through nanobody and protein A/G fusions preserve similar patterns of sequence discrimination. Together, these results establish RNA recording as a scalable approach to map sequence determinants of protein-RNA recognition across molecular and cellular contexts.

## Introduction

RNA-binding proteins (RBPs) play a central role in post-transcriptional gene regulation. They control RNA processing, nuclear export, translation, stability, and subcellular localization. RBPs also mediate assembly of larger ribonucleoprotein particles and granules, which play roles in diverse cellular functions. Dysregulated RBP-RNA interactions are implicated in a wide range of human diseases (Gebauer et al. 2021). A central goal is to identify which RNAs are bound by which RBPs and reveal the molecular bases of the interactions.

Powerful approaches exist to examine RBP-RNA interactions *in vitro* and in cellular contexts, but they have limitations. For example, systematic evolution of ligands by exponential enrichment (SELEX), Bind-N-Seq, Se-qRS, and RNA-MaP examine many thousands of RNA variants bound by an RBP *in vitro* (Tuerk and Gold 1990; Ellington and Szostak 1990; Lambert et al. 2014; Campbell et al. 2012; Becker et al. 2019; Buenrostro et al. 2014; Jarmoskaite et al. 2019). RBP variant libraries have also been interrogated through yeast three-hybrid and pooled biochemical assays (Zhou et al. 2021; Tasca et al. 2025). These strategies define consensus binding motifs across a range of affinities, but they lack human cellular context. Cellular immunoprecipitation-based approaches (e.g., RIP-seq) capture snapshots of RBPs bound to RNAs in more native contexts, but they also may capture non-native interactions that form in lysates (Zhao et al. 2010). Crosslinking strategies, such as CLIP-seq and its many derivatives, circumvent this limitation and provide nucleotide-resolution views of RBP binding in cells (Ule et al. 2005). However, these approaches are limited by crosslinking efficiency, antibody specificity, and biases introduced during crosslinking, RNA isolation, and sequencing. Thus, while current approaches have profoundly advanced our understanding of RBP-RNA interactions, it remains challenging to integrate *in vitro* mutational studies with *in vivo* profiling methods to achieve a more complete understanding of RBP function.

RNA recording-based approaches have emerged as a new strategy to uncover which RNAs are bound by an RBP in cells. In these approaches, an RBP of interest is fused to an RNA modification enzyme. The fusion protein covalently modifies bound RNAs, which can then be quantified by high-throughput sequencing. RNA modifying enzymes used for these studies include a poly(U) polymerase, engineered versions of the adenosine deaminase ADAR, and the cytosine deaminase APOBEC1 (Lapointe et al. 2015; McMahon et al. 2016; Rahman et al. 2018; Meyer 2019; Brannan et al. 2021). More recently, the *E. coli* adenosine deaminase TadA has been engineered to enhance its efficiency and allow it to modify a broad range of RNA substrates (Xiao et al. 2023; Lin et al. 2023). RBP target-mapping studies using deaminase fusions identified RNA targets that overlapped with those recovered by CLIP-based approaches (McMahon et al. 2016; Brannan et al. 2021; Lin et al. 2023). Moreover, the editing marks accumulate in the RNAs over time, require less input material, and can be multiplexed using orthogonal enzymes. Sequential recruitment of an RNA editor to an RBP using an antibody further enables profiling in fixed cells and tissues (Liang et al. 2024; Cheng et al. 2025). Thus, RNA recording provides a broader and complementary view of RBP-RNA interactions compared to direct binding-based approaches.

In this study, we extend RNA recording to scalable mutational studies of interactions between an RNA-binding protein and its RNA target. Specifically, we ask whether the extent of editing captures the molecular features required for an RBP to bind to its RNA target. Since RNA recording can be carried out with purified enzymes or in cells, it enables direct comparison of protein-RNA interactions between *in vitro* and cellular contexts. Further, recruitment of deaminases using antibody and nanobody fusions is a promising approach for applying RNA recording to a wide range of RBPs, but the specificity of these split recruitment strategies relative to a direct fusion remains unclear. Expression of RBP variant libraries in human cells could extend the same recording readout to determinants on the protein side. By testing the feasibility of these ideas using a model RBP-RNA system, here we demonstrate the utility of RNA recording for mapping the key molecular determinants of protein-RNA interactions at scale.

## Results

### Recruitment of a deaminase directs its editing to a target RNA

We first validated that fusing a deaminase to an RBP directs editing to a specific RNA *in vitro*. We selected TadA8.20, an evolved variant of the *E. coli* A-to-I deaminase TadA (Gaudelli et al. 2020) that efficiently edits RNA (Xiao et al. 2023). As the model RBP, we selected the 22 amino acid λN peptide, which specifically binds the boxB RNA stem-loop structure (Chattopadhyay et al. 1995). The λN–boxB system is widely used to tether proteins to RNAs for *in vivo* functional studies (De Gregorio et al. 1999; Baron-Benhamou et al. 2004). We used the 15-nucleotide *nutL* boxB stem loop (Tan and Frankel 1995). We purified TadA8.20 alone (hereafter, TadA) or as a TadA8.20-λN fusion protein (TadA-λN) (Figure 1A). We incubated the proteins with a model reporter RNA either with or without boxB (Figure 1B). Reporter RNAs then were reverse transcribed (RT), PCR amplified, and analyzed by long-read nanopore sequencing. With this strategy, A-to-I editing of RNA introduces A-to-G substitutions in individual sequenced DNA molecules. Hence, the frequency of A-to-G substitutions quantifies editing (Figure 1B).

**Figure 1:**
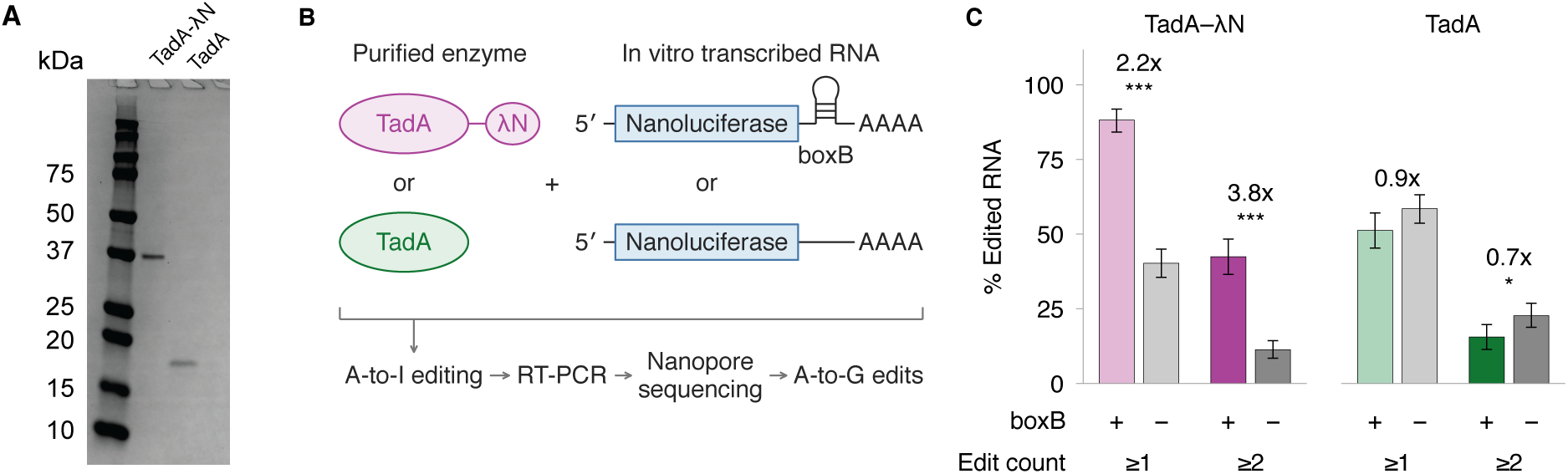
Recruitment of TadA increases editing of a boxB-containing reporter RNA. **(A)** SDS-PAGE analysis of purified TadA8.20–λN_1–22_ (TadA-λN) and TadA8.20 (TadA), visualized by Coomassie staining. **(B)** Reporter RNAs containing or lacking boxB were incubated with the indicated enzyme. A-to-I editing was detected as A-to-G substitutions by reverse transcription, PCR, and nanopore sequencing. **(C)** Bars show the percentage of indel-free reads containing at least one or at least two A-to-G substitutions for reporters containing or lacking boxB. Error bars indicate 95% confidence intervals from 100,000 binomial-bootstrap draws. Numbers above each pair are fold changes for +boxB relative to −boxB. *: P < 0.05, **: P < 0.01, ***: P < 0.001; no star indicates P ≥ 0.05 from two-sided Fisher exact tests on read counts.

Consistent with its high and non-specific activity (Gaudelli et al. 2020; Xiao et al. 2023), TadA alone introduced at least one edit into 50-60% of RNA molecules, regardless of whether they contained the boxB stem loop (Figure 1C). By contrast, TadA-λN edited boxB-containing RNAs 2.2-fold more than the RNA without boxB (∼90% versus ∼40%) (Figure 1C). Requiring two or more edits per molecule increased boxB-dependent editing enrichment to nearly 4-fold. Thus, the λN peptide directed the editor TadA to a specific RNA *in vitro*.

### A pooled assay captures target-specific editing by TadA

To examine RBP-RNA interactions at scale, we developed a pooled reporter assay with a deep sequencing readout of RNA editing. For these pooled assays, we extended boxB to 19 nucleotides by appending two G-C base pairs to its base (Legault et al. 1998). Each reporter contained this boxB sequence and a 17-nucleotide recorder with eight adenosines at positions 2, 4, 5, 8, 10, 13, 15, and 16 (Figure 2A) (Xiao et al. 2023). An adenosine-depleted spacer separated boxB from the recorder. Using pooled oligo synthesis, we systematically varied the boxB sequence, the recorder sequence, and the length of the spacer (0, 10, 20, or 30 nt) (Figure 2A). After *in vitro* transcription, we incubated the pooled RNA libraries with TadA-λN or TadA at sub-saturating (125 nM), equimolar (250 nM), or excess (500 nM) concentration relative to the total RNA pool (250 nM). We measured editing using RT-PCR followed by Illumina short read sequencing (Figure 2B). We compared reporters with cognate boxB (WT) or variants with a disrupted stem (MUT) that prevent boxB folding and λN binding (Figure 2A) (Chattopadhyay et al. 1995).

**Figure 2:**
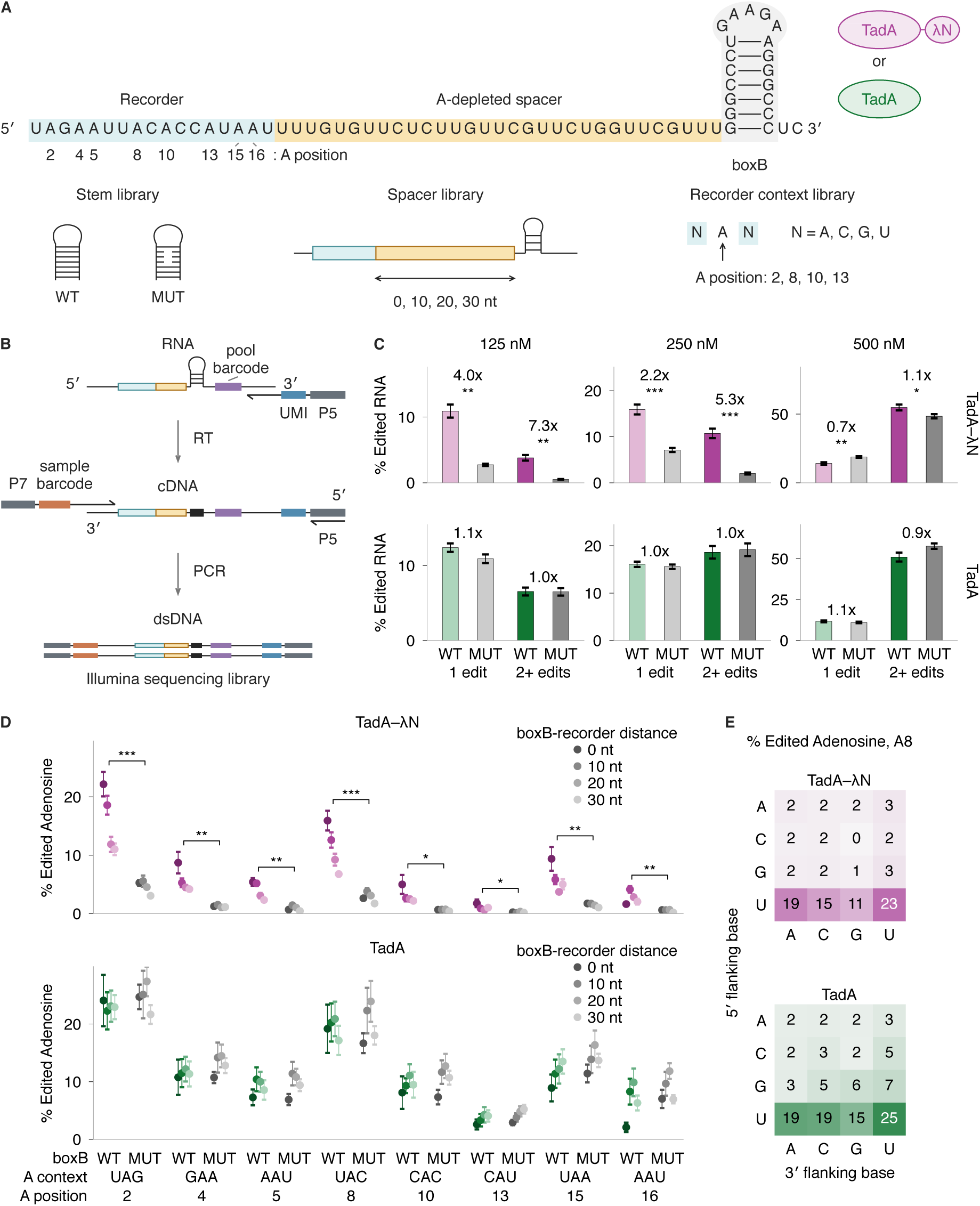
Pooled assay reveals boxB and sequence dependence of TadA-λN editing. **(A)** Reporter for the pooled sequencing assay, showing the recorder, spacer, and boxB. The stem library disrupts three boxB base pairs, the spacer library varies the boxB–recorder distance, and the recorder context library varies the nucleotides flanking adenosines 2, 8, 10, and 13. **(B)** Library preparation workflow for Illumina sequencing. The pool barcode distinguishes variants from different libraries; UMI, unique molecular identifier. **(C)** Percentage of unique RNA molecules with one edit or 2+ edits in the recorder region for WT or disrupted (MUT) boxB. RNA pool at 250 nM; enzyme concentration as indicated. Bars show means ± SEM across 22 WT constructs or 32 MUT constructs. Fold changes compare WT with MUT. **(D)** Percentage of molecules that carry an edit at each recorder adenosine, for TadA-λN and TadA, as a function of boxB–recorder distance. Points show means ± SEM across four to six WT or eight MUT constructs at each distance. **(E)** Percentage of molecules edited at recorder adenosine 8 by flanking context, pooled over all four boxB–recorder distances and 64 sequence backgrounds per distance. Color intensity varies in proportion to this percentage. D and E used 250 nM enzyme at 37 °C for 2 h. Brackets in D compare WT with MUT at each site; only significant comparisons are drawn. Tests use paired two-sided t-tests across eight matched window and placement groups. *: P < 0.05, **: P < 0.01, ***: P < 0.001; no star indicates P ≥ 0.05.

TadA-λN preferentially edited RNAs with the cognate boxB stem loop. When present at sub-saturating and equimolar concentrations, TadA-λN edited the recorder of wild-type boxB RNAs 2–4-fold more than MUT variants (Figure 2C). Specificity increased to more than 7-fold when considering only molecules with two or more edits at the sub-saturating concentration of TadA-λN. The relative enrichments persisted from 30 min to the longest incubation we tested (2 hours) (Supplementary Figure 1A). Consistent with saturation, this hallmark of specific binding disappeared when TadA-λN was in molar excess, with wild-type and MUT variants edited comparably or slightly favoring MUT (Figure 2C). TadA alone also lacked preference for RNAs with or without intact boxB at all concentrations, as expected. At sub-saturating and equimolar concentrations, TadA-λN editing of adenosines within boxB was comparable to the no-enzyme control, indicating that modification of boxB itself contributes minimally to the measured substitution effects (Supplementary Figure 1B). At excess enzyme concentration, wild-type boxB was protected from loop editing relative to stem-disrupted variants, consistent with protection by bound λN (Supplementary Figure 1B). Thus, our pooled assay detects preferential editing of RNAs that encode the cognate boxB site.

### TadA editing is sensitive to the context of the edited adenosine

Despite the directed evolution of engineered deaminases towards high activity and low sequence specificity, they often retain residual preference for editing nucleotides in particular contexts (Gaudelli et al. 2017; Gaudelli et al. 2020; Arbab et al. 2020; Xiao et al. 2024b; Shang et al. 2025). We therefore examined editing biases of TadA on our RNA reporter, which has eight adenosines located in different sequence contexts (Figure 2A). As in our aggregated analysis (Figure 2C), TadA-λN edited RNAs with its cognate boxB stem loop significantly more than those with a mutant version at all eight adenosine positions when present at sub-saturating and equimolar concentrations (Figure 2D and Supplementary Figure 2A). TadA alone lacked this preference at every position and concentration. Editing also declined modestly with distance, as recorders separated from boxB by 0, 10, 20, or 30 nt adenosine-depleted spacers showed progressively lower editing (Figure 2D). This decline required the boxB–λN interaction, since it was absent with mutant boxB or with TadA alone, and it weakened when TadA-λN was in molar excess (Supplementary Figure 2A,B). Across all RNAs and concentrations, both TadA-λN and TadA edited adenosines in UAG and UAC contexts more than the others (Figure 2D and Supplementary Figure 2A,B), with UAC being the context of the adenosine edited in tRNA by the ancestral wild-type *E. coli* enzyme (Wolf et al. 2002).

To assess these sequence context preferences more systematically, we examined TadA editing of a pooled RNA library with randomized sequence context around the recorder adenosines (Figure 2A). The randomization yielded four adenosines with all possible combinations of 5′ and 3′ flanking nucleotides. For both TadA and TadA-λN, adenosines with a 5′ uridine were edited 5–10-fold more than other sequence combinations (Figure 2E). Addition of a 3′ uridine (UAU) further enhanced editing by up to 2-fold, which yielded the highest percentage of edited adenosines across all contexts (Figure 2E and Supplementary Figure 3A). Conversely, adenosines that lacked U at the adjacent 5′ or 3′ nucleotide had the lowest percentage edited. We also examined published RNA sequencing from another TadA-derived RNA base editor, rABE, used to identify RBP binding sites *in vivo* (Lin et al. 2023). Among sites with detected editing in these datasets, mean editing was higher for adenosines flanked by a 5′ U or C (Supplementary Figure 3B). Collectively, our findings define intrinsic biases of the engineered TadA deaminase TadA8.20 in RNA recording experiments, with the highest editing occurring at UA dinucleotides.

### TadA-λN editing recovers sequence determinants of λN binding to boxB

Deaminase-RBP fusions in cells edit RNAs to variable extents (Medina-Munoz et al. 2024). Because endogenous RNAs differ in sequence, structure, abundance, and associated RBPs, whether the extent of editing reflects RBP binding strength remains unclear. We therefore used our *in vitro* pooled assay to examine how substitutions within boxB affect editing by TadA-λN. We designed six reporter libraries in which the boxB loop and stem regions were randomized in 3 or 4 nt windows, yielding 760 unique boxB variants (Figure 3A). To distinguish biological signal from statistical noise, we focused our analysis on the two recorder adenosines (A2 and A8) that had the highest percentage of edited molecules (Figure 2D). Editing at the two recorder sites was well correlated (r=0.71–0.99) across all boxB variants and all enzyme concentrations for both TadA-λN and TadA (Figure 3B, Supplementary Figure 4A,D). We therefore averaged the editing at these recorder sites for downstream analyses.

**Figure 3:**
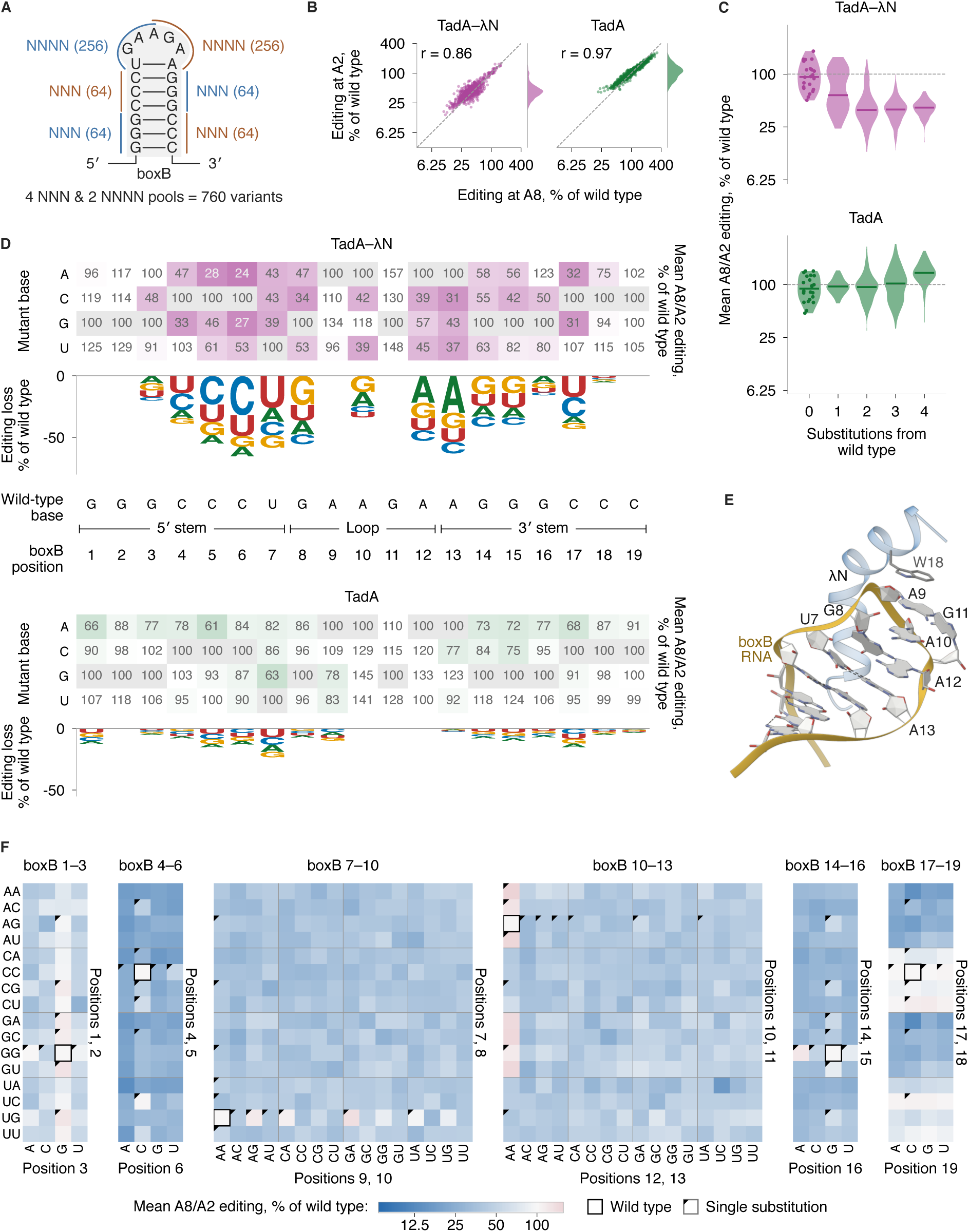
Mutational scanning of boxB recovers the determinants of λN binding. **(A)** boxB variant library. Six windows of three or four positions were randomized, giving 760 distinct variants. Designed sequences per window are in parentheses. **(B)** Editing at recorder adenosines A2 and A8 for the 759 variants other than wild type, each normalized to its matched wild type. Marginal densities show the A2 distribution, and the dashed line marks equal editing at the two sites. r is the Pearson correlation of log_2_ ratios. **(C)** Editing averaged across A8 and A2 as a function of the number of substitutions from wild-type boxB. Points at zero substitutions are the 22 wild-type constructs against the pooled wild-type value. Horizontal marks are medians. **(D)** Editing averaged across A8 and A2 of each single boxB mutant as a percentage of its matched wild type. Gray cells are the wild-type base. Logos show the mean loss across the three substitutions at each position, with letters scaled by the editing each base retains; positions without a net loss have no stack. **(E)** NMR structure of the λN-boxB complex (Legault et al. 1998), with the loop and closing base pair labeled. λN is blue and boxB is gold. **(F)** Editing averaged across A8 and A2 for every variant of all six randomized windows. B, C, and D compare TadA-λN with unfused TadA; F shows TadA-λN alone. All reactions used 250 nM enzyme and 250 nM RNA pool at 37 °C for 2 h.

We first analyzed the changes in editing aggregated across the boxB variant library. Mutations to boxB resulted in strikingly divergent effects on editing by TadA-λN and TadA. Median editing by TadA-λN across all boxB variants was reduced to around 40% of wild-type boxB when the enzyme was added at equimolar concentration (Figure 3B). By contrast, median editing by TadA increased slightly to ∼104% of wild-type boxB at the same concentration. This differential editing response between TadA-λN and TadA was accentuated at sub-saturating enzyme concentration and lost at molar excess (Supplementary Figure 4A,D). When the boxB variants were grouped by the number of substitutions away from the wild-type sequence, editing by TadA-λN progressively decreased up to two substitutions before leveling off (Figure 3C, Supplementary Figure 4B). This pattern was not seen at excess TadA-λN concentration relative to RNA or at any concentration of TadA (Figure 3C, Supplementary Figure 4B,E), but robust across different incubation times (Supplementary Figure 6B). Thus, editing by tethered TadA-λN is sensitive to substitutions away from the wild-type boxB sequence.

We then resolved the effect of substitutions at each position along boxB (Figure 3D, Supplementary Figure 4C, Supplementary Figure 5A). Within the boxB GAAGA pentaloop (positions 8–12), deleterious substitutions occurred at specific positions. Substitution of G_8_ or A_12_ to any other nucleotide strongly reduced editing by TadA-λN (Figure 3D, 3F). A_10_ tolerated the other purine but not a pyrimidine (Figure 3D, Supplementary Figure 4C). These mutational effects reflect the known architecture of λN bound to the GNRNA loop (Figure 3E), which is a variant of the common GNRA tetraloop (Thapar et al. 2014). λN contacts the G_8_ nucleobase directly via Gln4 and Arg7 (Legault et al. 1998; Schärpf et al. 2000). While not contacted directly by λN, A_12_ forms a sheared base pair with G_8_ required for the boxB loop to fold properly (Legault et al. 1998; Schärpf et al. 2000). The purine rings of A_10_ stack in between A_9_ and A_12_, which substitution by G is not expected to disrupt. Accordingly, disruptive substitutions at boxB positions A_10_ and A_12_ impair λN binding *in vitro* (Chattopadhyay et al. 1995; Tan and Frankel 1995; Su et al. 1997b). While A_9_ also contacts λN, the interaction occurs via pi-stacking with Trp18 and contacts with its ribose ring (Legault et al. 1998; Schärpf et al. 2000; Xia et al. 2003). All three substitutions at A_9_ retained near-wild-type editing (Figure 3D,F, Supplementary Figure 4C). G_11_ extrudes from the folded loop, lacks contacts with λN (Legault et al. 1998; Schärpf et al. 2000), and enables a downstream transcriptional response *in vivo* (Mogridge et al. 1995; Legault et al. 1998). Accordingly, substitutions at G_11_ yielded wild-type levels of editing (Figure 3D). Importantly, the above disruptive substitutions were strongly attenuated with TadA alone or molar excess TadA-λN (Figure 3D, Supplementary Figure 4F, Supplementary Figure 5B).

Deleterious substitutions also clustered asymmetrically within the boxB stem. Along the 5′ strand, substitution of C_5_, C_6_, and U_7_ strongly reduced editing by TadA-λN (Figure 3D,F, Supplementary Figure 4C). A similar reduction occurred at A_13_, the 3′ partner of U_7_. However, reductions at the remaining 3′ partners were much more modest; G_15_ and G_14_ variants retained about half of wild type editing levels. These findings again reflect the architecture of the λN–boxB complex. λN contacts the phosphate backbone of the 5′ strand via its conserved arginine-rich motif (Arg6, Arg7, Arg8, Arg10, Arg11) and packs within the major groove near C_4_ and C_5_ via Ala3 (Legault et al. 1998; Schärpf et al. 2000). Each of these contacts contributes to high-affinity binding (Cilley and Williamson 1997; Su et al. 1997b). Furthermore, the closing U_7_-A_13_ base pair is a primary determinant of λN specificity (Legault et al. 1998), and its inversion reduces λN binding more than 20-fold (Tan and Frankel 1995; Legault et al. 1998). By contrast, λN lacks direct contacts with the rest of the 3′ strand, and the modest effects on editing likely reflect loss of base pairing. Purine substitutions at C_4_ and C_17_ reduced editing, whereas U was tolerated at both positions (Figure 3D, Supplementary Figure 4C), consistent with tolerance for wobble (G-U) pairs at these positions. Finally, the outermost two appended base pairs tolerated most substitutions (Figure 3D, 3F), suggesting that pairing at these positions is dispensable for λN binding.

Collectively, our findings demonstrate that TadA-λN provides a binding-dependent editing readout *in vitro*. After a single pooled incubation, RNA recording recovered key sequence determinants of λN-boxB recognition established by prior biochemical, genetic, and structural studies.

### Split recruitment of deaminase maintains λN-boxB specificity

Direct fusions of deaminases to RBPs require prior genetic engineering of each RBP. Recent studies have instead recruited RNA editors through antibodies to profile RBP-associated RNAs in fixed cells and tissues (Liang et al. 2024; Cheng et al. 2025). However, the specificity of split recruitment relative to direct RBP fusions remains unclear. To address this gap, we compared direct editing by TadA-λN to two split recruitment strategies using our pooled *in vitro* assay (Figure 4A). For both, we purified λN fused to GFP (λN-EGFP) to serve as the model RBP (Supplementary Figure 7A). We then purified TadA fused to a GFP nanobody (TadA-GFPNb), which recognizes λN-EGFP directly, as well as TadA fused to protein A/G (pAG-TadA), which recognizes an anti-GFP antibody bound to λN-EGFP (Supplementary Figure 7A).

**Figure 4:**
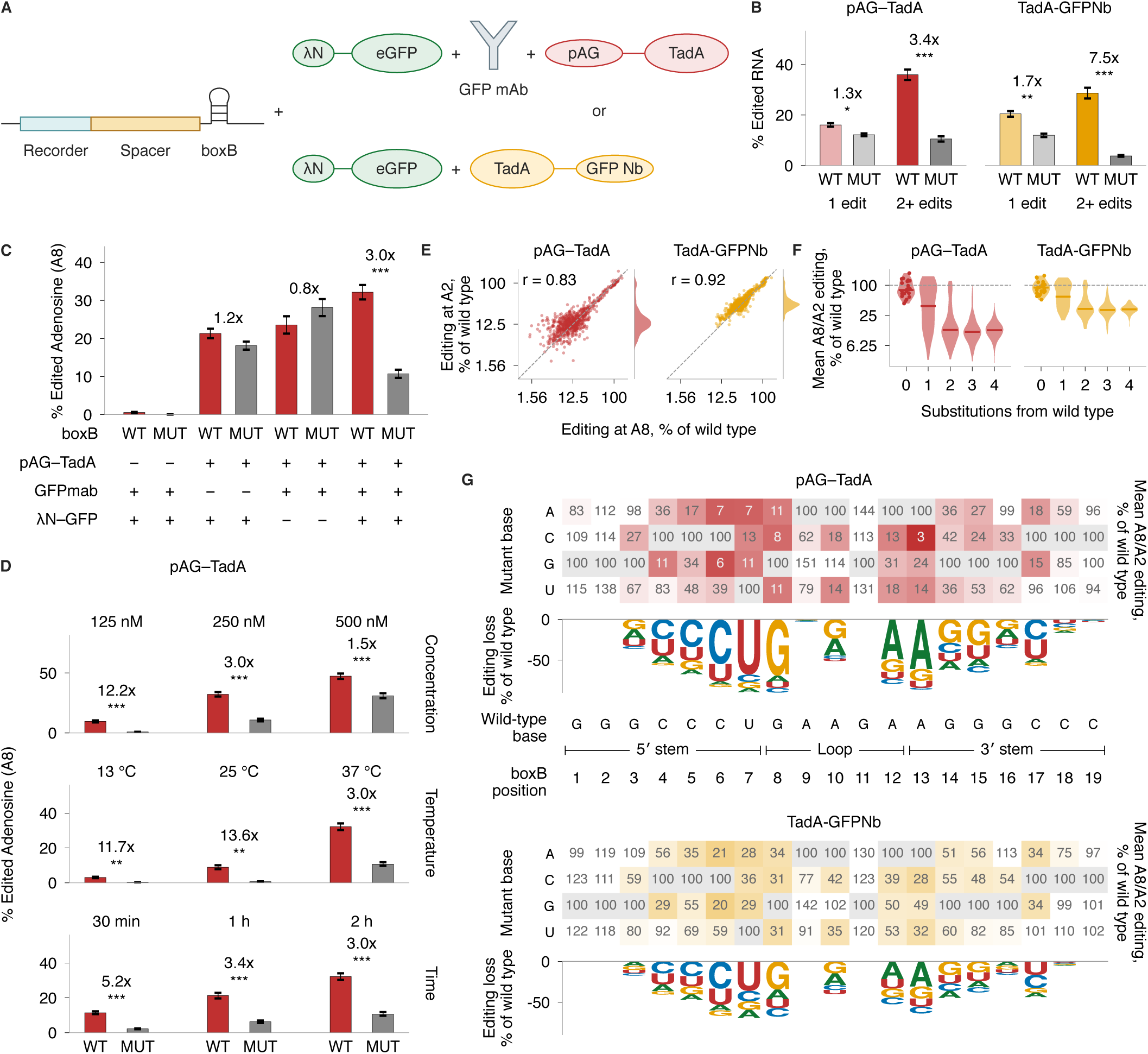
Split recruitment of TadA preserves editing specificity and boxB sequence determinants. **(A)** Recruitment of TadA to boxB reporters through λN-EGFP, using either an anti-GFP monoclonal antibody with the protein A/G fusion pAG-TadA or the GFP nanobody fusion TadA-GFPNb. **(B)** Percentage of unique RNA molecules with one edit or 2+ edits in the recorder region for WT or stem-disrupted (MUT) boxB. **(C)** Editing at recorder adenosine A8 with the indicated components omitted. **(D)** A8 editing by pAG-TadA as a function of concentration, temperature, and incubation time. Concentration applies to all recruitment components together. Bars in B-D show means ± SEM across 22 WT or 32 MUT constructs, as in Figure 2C. Fold changes compare WT with MUT (paired two-sided t-tests). **(E)** Editing at A2 and A8 for the 759 variants other than wild type, as in Figure 3B. Three pAG-TadA variants with no edited molecule are not shown and are excluded from r. **(F)** Editing averaged across A8 and A2 as a function of the number of substitutions from wild-type boxB, as in Figure 3C. **(G)** Editing averaged across A8 and A2 of each single boxB mutant, as in Figure 3D. B and C used 250 nM of each component, and D varies one condition at a time from that shared reaction. E-G used 125 nM pAG-TadA or 250 nM TadA-GFPNb. All reactions were at 37 °C for 2 h except where indicated in D. *: P < 0.05, **: P < 0.01, ***: P < 0.001; no star indicates P ≥ 0.05.

With all components present at equimolar concentrations (250 nM), TadA-GFPNb and pAG-TadA both showed higher editing of RNAs with intact boxB relative to stem-disrupted variants (Figure 4B, Supplementary Figure 7C,D). Wild-type-to-mutant editing enrichment was 3–8-fold when we quantified multiple edits (Figure 4B) or edits at the two recorder positions A2 and A8 with the highest percentage of edited molecules (Supplementary Figure 7C,D). Using the more complex pAG-TadA system, we investigated the component and reaction conditions driving specific editing of wild-type boxB over stem-disrupted variants. Omission of either the EGFP antibody or λN-EGFP prevented specific editing of wild-type boxB RNA by pAG-TadA, whereas pAG-TadA omission eliminated editing altogether (Figure 4C and Supplementary Figure 7B). The same pattern held across whole RNA molecules with single or multiple edits, and specificity increased to 16-fold for multiply edited RNAs at 125 nM of each component (Supplementary Figure 7F,G). Specificity also inversely correlated with editing activity; the highest specificity occurred in conditions where the fewest RNA molecules were edited, including lower enzyme concentration, lower temperature, and shorter incubation times (Figure 4D, Supplementary Figure 7E). These findings indicate that split recruitment of TadA can be controlled to balance activity with specificity.

Split recruitment of the deaminase also recovered the same boxB sequence determinants as the direct fusion when tested against the boxB variant library (Figure 3A). For both nanobody-and antibody-driven recruitment, editing at the recorder adenosines A2 and A8 correlated strongly (r = 0.92 and 0.83) across all boxB library variants (Figure 4E), and therefore, we averaged across their editing percentages for further analyses. Direct comparison of matched boxB variants at 250 nM of each component showed correlated effects between direct and split recruitment (r = 0.76 for pAG-TadA and 0.89 for TadA-GFPNb; Supplementary Figure 7H). Editing progressively decreased as the number of substitutions away from wild-type boxB increased (Figure 4F), similar to the direct fusion (Figure 3C). Position-resolved analyses of editing by the split editors recovered the known architecture of the λN-boxB complex. In the GNRNA pentaloop (boxB positions 8–12), G_8_ and A_12_ substitutions strongly diminished editing, and A_10_ tolerated the other purine but not a pyrimidine (Figure 4G). Both A_9_ and G_11_ tolerated all substitutions. Within the stem, substitutions at the loop-proximal U_7_ and its partner A_13_ strongly reduced editing. Elsewhere, effects of stem disruptions progressively decreased away from the loop (Figure 4G). Disruptions in the 5′ strand of the boxB stem had a stronger effect on editing than on the 3′ strand, similar to our earlier observations with the direct fusion. We also observed stronger effects of purine substitutions at C_4_ and C_17_ as before.

Together, our findings demonstrate that split recruitment preserves sequence discrimination across the boxB library. The tradeoff between editing specificity and activity of the split strategy can be tuned by changes in concentration, temperature, and incubation time, enabling versatile future applications.

### TadA-λN editing in cells recovers boxB recognition determinants

We next asked whether editing patterns observed in vitro were preserved in human cells. We took advantage of our ability to integrate both the editing enzyme and variant boxB libraries into the *AAVS1* locus of HEK293T cells using our previously developed landing pad system for Bxb1 recombination (Nugent et al. 2024). Using this pooled strategy, an EGFP reporter with a single boxB variant and the recorder embedded in the 3′UTR and the TadA-λN enzyme are expressed off separate doxycycline-regulated promoters within each cell (Figure 5A). After doxycycline induction for 72 hours, we isolated RNA and sequenced the recorder region as before (Figure 2B). We detected all 760 boxB variants, with coverage varying across two orders of magnitude (Figure 5B).

**Figure 5:**
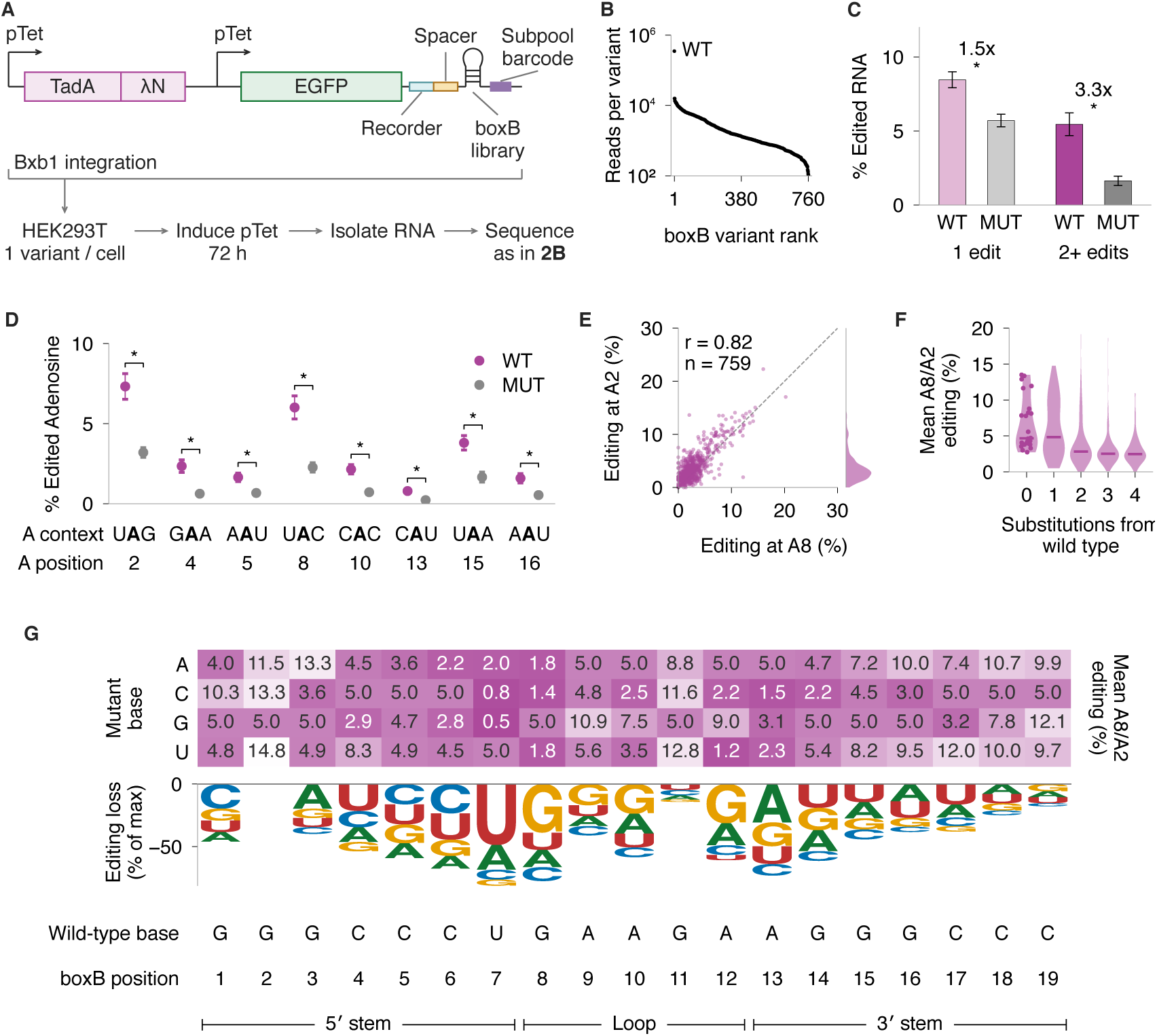
TadA-λN editing in human cells recapitulates boxB sequence determinants. **(A)** boxB variant library and TadA-λN expressed from the *AAVS1* genomic locus in HEK293T cells. Bxb1 recombination gives a single variant per cell; libraries were induced with doxycycline for 72 h and sequenced as in Figure 2B. **(B)** Read coverage of the 760 boxB variants, ranked. **(C)** Percentage of unique RNA molecules with one edit or 2+ edits in the recorder region for WT or stem-disrupted (MUT) boxB. Bars show means ± SEM across 24 WT or 32 MUT constructs. Fold changes compare WT with MUT (paired two-sided t-tests). **(D)** Editing at each recorder adenosine and its sequence context. Error bars are SEM across constructs. Brackets compare WT with MUT at each site, across eight matched barcode groups; only significant comparisons are drawn. **(E)** Editing at A2 and A8 for the 759 designed boxB mutants, with sequences unedited at a site drawn at zero on that axis. r is the Pearson correlation. **(F)** Editing averaged across A8 and A2 as a function of the number of substitutions from wild-type boxB, and includes 57, 216, 324, and 162 variants for one to four substitutions, respectively. Points at zero substitutions are the 24 wild-type constructs. Horizontal marks are medians. **(G)** Editing averaged across A8 and A2 for each single boxB mutant. Colors show the unnormalized editing percentage. The logo shows each position’s reduction from the highest position mean, with letters scaled by the editing each base retains. *: P < 0.05.

TadA-λN preferentially edited RNAs with wild-type boxB over stem-disrupted variants (Figure 5C), as observed in vitro (Figure 2C). This enrichment increased to more than 3-fold on RNAs with multiple edits. As expected, we did not detect editing of the recorder in cells without TadA-λN (Supplementary Figure 8A). Specific editing of RNAs with wild-type boxB extended across all eight adenosines in the recorder, with UA contexts again being the most edited (Figure 5D). Editing at positions 2 (UAG) and 8 (UAC) also remained correlated across all 759 designed boxB variants (r = 0.82) (Figure 5E), and we therefore averaged across these positions for further analyses. The fraction of wild-type boxB molecules with exactly one recorder edit was similar in cells and at 125 nM enzyme in vitro (13.9% versus 14.7%; Figure 2C), although wild-type-to-mutant enrichment was 2-to 3-fold lower because stem-disrupted RNAs were edited about twice as much in cells. Single substitutions in boxB yielded a range of editing activities, and multiple substitutions strongly reduced editing (Figure 5F).

TadA-λN expressed in cells recovered key determinants of λN–boxB recognition. In the boxB GNRNA pentaloop, G_8_ substitutions strongly diminished editing, A_10_ and A_12_ tolerated the other purine but not a pyrimidine, A_9_ modestly reduced editing, and G_11_ again proved flexible (Figure 5G). Within the stem, substitutions at the loop-proximal U_7_ and A_13_ also strongly reduced editing. The largest effects persisted on the 5′strand of the stem with direct λN contacts, and progressively decreased at distal stem locations away from the loop. The strongest mutational effects observed *in vitro* (Figure 3D) were generally conserved in cells, whereas smaller effects were attenuated. Editing effects across boxB variants showed a moderate correlation between in vitro and in vivo measurements (r = 0.48; Supplementary Figure 8B). Agreement was stronger for single substitutions (r = 0.78) and stem-window variants (r = 0.76) than for loop-window variants (r = 0.37). Most loop variants reduced in vitro editing to near-baseline levels, limiting the range over which their effects could be resolved. Overall, the same recorder readout captures boxB sequence requirements in biochemical reactions and human cells.

### Mutational scanning of λN identifies residues required for boxB binding

Protein-side mutational studies have used individual purified variants (Jessen et al. 1991; Johansson et al. 1998; Mikulecky et al. 2004; Mercante et al. 2006), transcription-dependent yeast selections (Zhou et al. 2021), and, more recently, pooled biochemical assays with peptide barcodes and mass spectrometry (Tasca et al. 2025). We reasoned that combining in vivo RNA recording with our earlier RNA-linked perturbation strategy (Nugent et al. 2024) could provide a scalable approach for mutational studies of RBP binding. To test this idea, we performed single-codon saturation mutagenesis at each of the 22 λN positions, giving 1,387 unique codon variants encoding 441 distinct protein sequences (Figure 6A). Each λN variant was fused to TadA and expressed in HEK293T cells using Bxb1 recombination as before. We co-expressed an EGFP reporter with a 3′UTR that contained wild-type boxB, the A-rich recorder region, and a 20-nucleotide barcode linked to a unique λN variant (Nugent et al. 2024).

**Figure 6:**
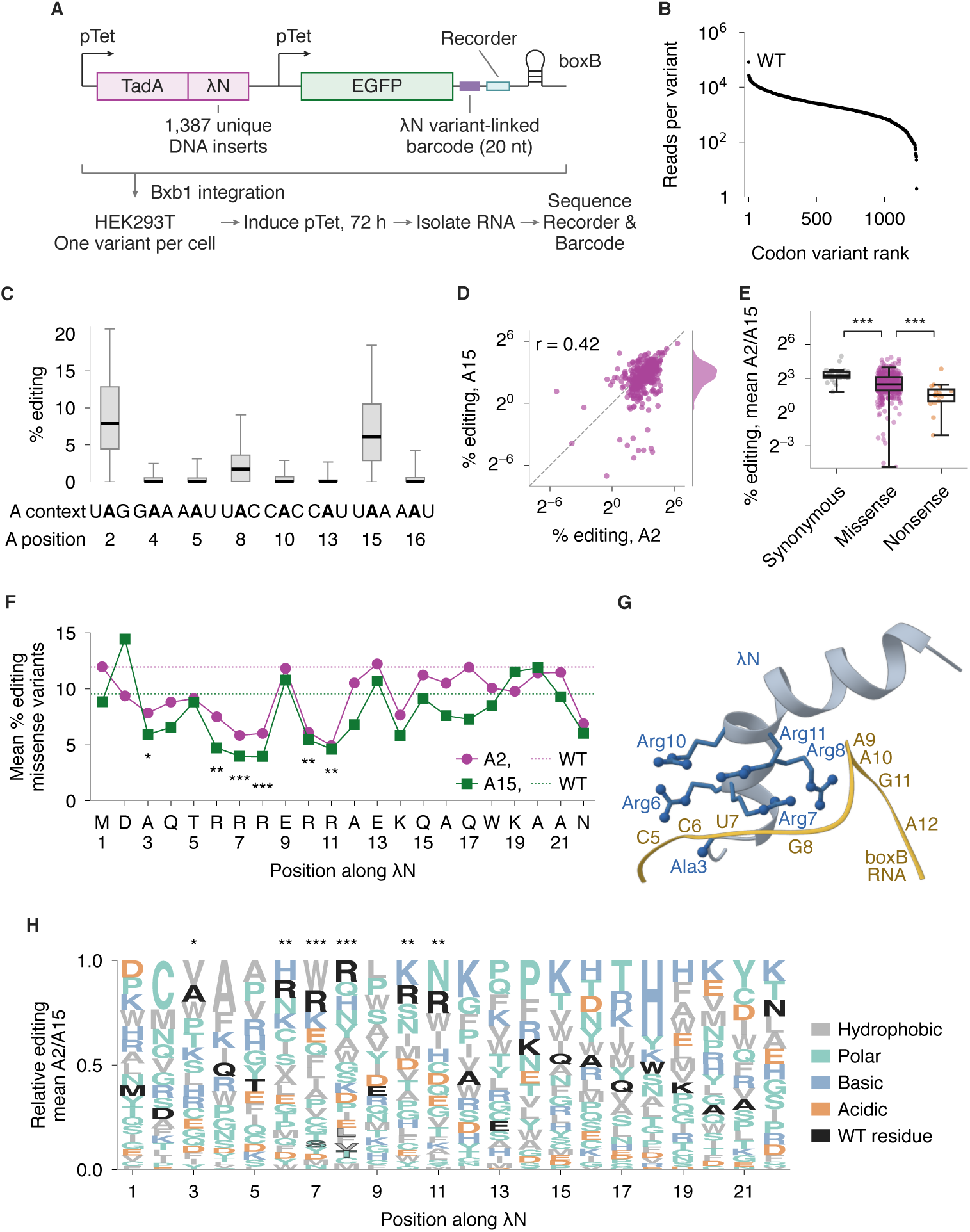
Deep mutational scanning identifies λN residues required for boxB binding. **(A)** λN single-codon saturation library (1,408 position-codon designs; 1,387 unique DNA inserts) fused to TadA and expressed with a boxB reporter whose 3′ UTR carries a 20-nt barcode linked to each λN codon variant. The library was integrated into the *AAVS1* genomic locus of HEK293T cells, induced with doxycycline for 72 h, and sequenced across the recorder and barcode. **(B)** Read coverage of the 1,236 detected codon variants, ranked. **(C)** Editing at each recorder adenosine and its sequence context across 436 amino acid variants. Boxes span quartiles with the median, and whiskers the 5th–95th percentiles. **(D)** Editing at A2 and A15 for the 381 amino acid variants with editing detected at both sites. r is the Pearson correlation of log_2_ rates. **(E)** Editing averaged across A2 and A15 for codon variants with at least five barcodes at each site. Boxes span quartiles with the median, and whiskers extend to 1.5 interquartile ranges. Brackets compare neighboring classes (two-sided Mann–Whitney tests). **(F)** Mean editing of missense variants at each λN position, with A2 and A15 shown separately. Dotted lines mark the pooled synonymous wild type for each site. **(G)** NMR structure of the λN-boxB complex (Legault et al. 1998), with the six positions marked in F highlighted in blue. **(H)** Relative editing of each amino acid within a position, from the mean of A2 and A15. Letters are ordered by increasing editing from bottom to top, and the wild-type residue is black. Outlined letters mark substitutions that differ from pooled synonymous wild type at both sites (BH-adjusted P < 0.05 at each). Stars in F and H compare position means with pooled synonymous wild type, using the larger site-specific BH-adjusted P value (two-sided t tests). *: adjusted P < 0.05, **: adjusted P < 0.01, ***: adjusted P < 0.001.

After doxycycline induction for 72 hours, we isolated RNA and sequenced the recorder region and barcode as before (Figure 2B). We recovered 1,236 codon λN variants after sequencing, which represented 436 distinct λN protein variants (Figure 6B).

We first analyzed editing within the recorder aggregated across the λN library. Recorder positions with a UA dinucleotide context (2, 8, 15) showed the highest editing (Figure 6C). The decreased editing at position 8 likely reflects sequence beyond its directly adjacent nucleotides, which differs in this reporter (Figure 6A). We therefore quantified effects at positions 2 (UAG) and 15 (UAA). Editing at these positions correlated across variants, although less tightly than in boxB libraries (r = 0.42) (Figure 6D). For the codon-class comparison, we required at least five distinct barcodes at both positions, retaining 394 of the 1,236 detected variants (Figure 6E). As expected, synonymous codon variants had the highest editing and nonsense variants the lowest, with missense variants spanning from wild-type to background levels of editing (median editing of 9.6% for synonymous, 5.5% for missense, and 2.9% for nonsense variants) (Figure 6E). The nonsense variants provide internal controls for reduced recording; their stop codons truncate λN downstream of the TadA coding sequence.

We next examined the effect of individual variants at each position of λN on editing. Variants of Ala3, Arg6, Arg7, Arg8, Arg10, and Arg11 decreased editing to the lowest levels along the entire λN peptide, and were the only six positions that differed significantly from the pooled synonymous wild type at both recorder positions (Figure 6F). Arginine ranked at or near the top of residues at positions 6, 7, 8, 10, and 11, while alanine and other small residues ranked among the highest at position 3 (Figure 6H). These requirements match the architecture of the λN-boxB complex (Legault et al. 1998; Schärpf et al. 2000) (Figure 6G). The arginines contact the phosphate backbone of the stem or make base-and architecture-specific contacts in the loop and its closing base pair (Su et al. 1997a; Legault et al. 1998), while the Ala3 methyl group packs within the major groove of the stem between C_4_ and C_5_ (Legault et al. 1998). These same arginine residues and Ala3 underlie both λN binding *in vitro* and its regulatory activity *in vivo* (Franklin 1993; Su et al. 1997a; Austin et al. 2003). Gln4, Lys14, and Trp18 contact boxB in structures of the complex (Su et al. 1997a; Legault et al. 1998; Schärpf et al. 2000), yet their position-averaged effects did not reach significance (Figure 6F). Su et al. found that all 19 substitutions at both Ala3 and Trp18 impaired peptide binding; our scan recovered the Ala3 requirement but did not resolve the established contribution of Trp18 (Su et al. 1997a). Since RNA editing also depends on fusion abundance and activity, the substitution effects identified in our assay might not mirror binding measurements with purified peptides.

Collectively, our mutational analyses from the RBP perspective using RNA recording in human cells recapitulated the architecture of the λN-boxB complex. Importantly, our RNA and protein mutational findings converged. The same arginine-rich motif identified as critical above engages the 5′ strand of the boxB stem and the G_8_ nucleobase (Figures 6F-G), and substitutions within boxB at those positions reduced editing most severely on the RNA side (Figure 3D). Our findings demonstrate that RNA recording recovers convergent sequence requirements on the RNA and protein sides of the λN-boxB interaction.

## Discussion

Here, we present a high-throughput strategy to interrogate the molecular interactions that underlie protein– RNA binding. Our RNA recording strategy builds on RNA editing approaches commonly used to map transcriptome-wide RBP binding sites (McMahon et al. 2016; Brannan et al. 2021; Lin et al. 2023). In parallel with our study, pooled RNA libraries have been read out in cells both by ADAR-mediated recording with a quantitative dose-response framework (Kretov et al. 2026) and by structural probing that quantifies protein occupancy from RNA hairpin reactivity (Thurm et al. 2026). RNA recording provides a common readout for mapping sequence requirements on both sides of a protein-RNA interaction, comparing biochemical and cellular settings, and testing direct versus split recruitment. By combining RNA editing with pooled variant libraries, we recovered convergent RNA and protein determinants of λN-boxB recognition. The strongest RNA mutational effects were generally conserved in cells, whereas smaller effects were attenuated. Our split recruitment strategy preserved mutational discrimination, supporting its application to mutational profiling alongside antibody-directed and chemically induced RNA-recording approaches (Liang et al. 2024; Cheng et al. 2025; Seo and Kleiner 2023). Together, the strategies described here offer complementary ways to tune RNA recording for scalable studies of protein-RNA recognition.

## Author Contributions

R.A.B. designed research, performed experiments, analyzed data, and wrote the first draft of the manuscript. H.P. and O.N. performed experiments. Y.X. and W.T. contributed reagents and technical expertise. A.R.S. and C.P.L. designed research, analyzed data, wrote the manuscript, supervised the project, and acquired funding.

## Acknowledgements

We thank members of the Subramaniam and Lapointe labs, the Basic Sciences Division, and the Computational Biology Program at Fred Hutch for assistance with the project and discussions and feedback on the manuscript. We thank Tao Pan for suggesting the use of TadA8.20 for RNA recording. This research was funded by NIH R35 GM119835 (A.R.S.), NIH R01 AT012826 (A.R.S.), NSF MCB 1846521 (A.R.S.), NSF PRFB 2209388 (R.A.B.), NIH R00 GM144678 (C.P.L.), NIH R35 GM160398 (C.P.L.), and the Damon Runyon Cancer Research Foundation DFS-49-22 (C.P.L.). This research was supported by the Genomics, Flow Cytometry, and Proteomics Shared Resources of the Fred Hutch/University of Washington Cancer Consortium (NIH P30 CA015704) and Fred Hutch Scientific Computing (NIH S10 OD020069 and NIH S10 OD028685). The funders had no role in study design, data collection and analysis, decision to publish, or preparation of the manuscript.

## Data and Code Availability

Illumina sequencing data generated in this study are available at the NCBI Sequence Read Archive (SRA) under BioProject PRJNA1255650, with per-run accessions listed in the analysis repository. All other data and programming code associated with this manuscript are publicly available at https://github.com/rasilab/tada_boxb, including the source data behind every figure panel as a separate CSV file.

## Materials and Methods

Plasmids, oligonucleotides, and cell lines used in this study are listed in Supplementary Tables S1-S3, and the designed oligonucleotide pools in Supplementary Tables S4-S6. DNA sequences of plasmids used in this study are available at https://github.com/rasilab/tada_boxb/. Information not included below can be requested at https://github.com/rasilab/tada_boxb/issues/.

## Plasmid construction

### Protein expression plasmids

The parent TadA8.20 expression plasmid in a pET28a backbone was described previously (Xiao et al. 2023). To generate TadA-GFPNb expression vector (pAS95), Gibson assembly was performed using this plasmid as a backbone along with the following components: a 14xHis-Avi-SumoEu1 fragment (amplified with primers oRB96 and oRB97), a 48-amino-acid extended XTEN linker (Yarnall et al. 2023), and the GFPNb sequence. This vector (pAS95) served as the template for constructing additional protein expression plasmids.

To generate the TadA-λN expression plasmid (pAS335), pAS95 was digested with BamHI and XhoI to remove the TadA-GFPNb region. The resulting backbone was assembled via Gibson with a synthetic TadA-λN fragment (oAS2160, ordered from GenScript).

To construct the pAG-TadA expression plasmid (pAS428), pAS95 was digested with XhoI and SacI to remove the XTEN-GFPNb region. The backbone was assembled with the pAG sequence (amplified from the pAG/MNase plasmid, Addgene #123461, using primers oRB226, oRB227, and oRB228; a gift from the Henikoff lab) and a pAG-specific linker (amplified from pAG/MNase using primers oRB231 and oRB242).

To generate the λN-EGFP expression plasmid (pAS334), pAS95 was digested with BamHI and XhoI to remove the TadA-GFPNb region. The resulting backbone was assembled via Gibson with a synthetic λN-EGFP fragment (oAS2159, ordered from GenScript).

### Reporter libraries for *in vivo* expression

First, the TadA-λN coding sequence was amplified from its expression plasmid (pAS335) using primers oRB247/oRB248. This fragment was cloned using Gibson assembly into a backbone vector containing a cHS4 insulator sequence and pTet Doxycycline inducible promoter sequence, yielding pAS441.

Next, pAS441 was digested with MluI and AgeI and assembled with two PCR fragments by Gibson assembly. EGFP was amplified from pAS332 using oRB257/oRB255, and the rbGlobin_pA terminator and pTet promoter were amplified from pAS155 using oRB258/oPB496. This assembly yielded pAS444 (TadA-λN + EGFP). This promoter-EGFP plasmid was then digested with NotI and assembled by Gibson with an EcoRV-digested parent vector (pAS457) containing attB sequences for Bxb1 recombinase integration into the genome, mCherry, and puromycin resistance as markers for integration, yielding pAS473. Following this, pAS473 was digested with NotI, and the boxB reporter library oligo pool (oRB262) was inserted via Gibson assembly to create pAS476. The pAS476 library was cloned in NEB10-beta cells with 0.2-0.3 million colonies to retain library complexity. The noeditor library pAS507 was generated from pAS476 by removing the pTet-TadA-λN cassette with EcoRV and MluI and closing the backbone with annealed oligos oRB285/oRB286, giving 0.3-0.4 million colonies. This retained the inducible EGFP-boxB reporter library.

To construct the λN site saturation mutagenesis library, pAS157 was digested with MluI and KpnI. The backbone was assembled by Gibson with a TadA-XTEN fragment amplified from the reporter intermediate pAS472 using oRB276/oRB283, yielding pAS493. In parallel, the boxB reporter sequence was amplified with oRB268/oRB269 and cloned into a separate plasmid bearing the attB site using the NEBuilder HiFi DNA Assembly system (NEB), yielding pAS495. pAS493 was then digested with NotI and pAS495 with EcoRV, and the two were assembled via Gibson to generate pAS496.

The λN site saturation mutagenesis library was synthesized as an oligo pool (oRB275) by IDT. This pool was amplified using primers oRB271 and oRB272 to append a unique 20-nt DNA barcode and homology arms for Gibson assembly. The plasmid pAS496 was digested with AgeI and MluI, and the barcoded λN mutagenesis library was inserted in-frame with the upstream TadA-XTEN via Gibson assembly. The final transformants were bottlenecked to 17,500 colonies, providing >10× coverage of the 1,387 unique λN DNA inserts in the library. The resulting plasmid pool (pAS499) was sequenced to link each 20-nt barcode to its corresponding λN variant.

To complete the functional reporter construct, pAS499 was digested with MluI and AgeI, and a fragment containing the rbGlobin terminator, pTet promoter, and EGFP coding sequence (from pAS444 digested with the same two enzymes) was inserted by ligation. This positioned the barcode and boxB reporter within the 3′ UTR of EGFP. This assembly yielded pAS506 at approximately 3 million colonies, but roughly 90% of the pool was uncut parent plasmid. The correct 7,760 bp species was therefore recovered by digesting pAS506 with BglII and NheI, gel-purifying that band, and religating it, which gave approximately 2 million colonies and maintained high representation of λN variants and barcodes. The resulting plasmid pool (pAS517) was used for genomic integration into cell lines.

### Protein purification

#### λN-EGFP

The plasmid was transformed into Rosetta 2 cells purchased from the UC Berkeley QB3 MacroLab and grown overnight at 37 °C on LB agar plates supplemented with 50 µg/mL of kanamycin. Liquid cultures of single colonies were grown at 37 °C in LB supplemented with kanamycin. At an OD_600_ of 0.5, isopropyl β-D-1thiogalactopyranoside (IPTG) was added at a final concentration of 0.5 mM. After two hours at 37 °C, the cultures were shifted to 16 °C and grown overnight. Cells were harvested by centrifugation at 5,000 x g for 10 minutes at 4 °C in a Fiberlite F9 rotor (ThermoFisher, cat # 096-061075). Cells were lysed by sonication in lysis buffer (20 mM Tris-HCl pH 8.0, 300 mM NaCl, 10% (v/v) glycerol, 30 mM imidazole, and 5 mM β-mercaptoethanol) supplemented with protease inhibitors (ThermoFisher, cat # A32965). Lysates were cleared by centrifugation at 27,000 x g for 45 minutes at 4 °C in a Sorvall SS-34 rotor. Clarified lysate was loaded onto Ni-NTA resin (Qiagen, cat # 30210) equilibrated in lysis buffer in a gravity flow column. The resin was then washed with 10 column volumes (CV) of lysis buffer, 10 CV of wash buffer (20 mM Tris-HCl pH 8.0, 1000 mM NaCl, 10% (v/v) glycerol, 30 mM imidazole, and 5 mM β-mercaptoethanol), and 10 CV of lysis buffer. Recombinant protein was eluted in five CV of elution buffer (20 mM Tris-HCl pH 8.0, 300 mM NaCl, 10% (v/v) glycerol, 300 mM imidazole, and 5 mM β-mercaptoethanol). Fractions with recombinant protein were identified by SDS-PAGE analysis. Relevant fractions were dialyzed overnight at 4 °C into dialysis buffer (20 mM Tris-HCl pH 8.0, 250 mM NaCl, 10% (v/v) glycerol, and 5 mM β-mercaptoethanol) in the presence of TEV protease. TEV protease and the cleaved 14xHis-AviSumoEu1 tag were captured via a subtractive Ni-NTA gravity column equilibrated in dialysis buffer supplemented with 30 mM imidazole. The flowthrough was collected and concentrated to 500 µL with a 10k MWCO concentrator (MilliporeSigma, cat # UFC901096) then applied to a 24 mL Superdex 200 Increase 10/300 GL SEC column (Cytiva, cat # 28990944). Fractions that contained purified λN-EGFP were concentrated, frozen in liquid N2, and stored at –80 °C.

#### TadA-GFPNb

Protein was purified as described above with the following modifications. Following the subtractive Ni-NTA step, the protein was diluted to 75 mM NaCl using 20 mM Tris-HCl pH 7.5 and applied to a 1 mL heparin column (Cytiva, cat # 17040601). Purified protein was eluted using a 50 to 1000 mM NaCl gradient in the absence of reducing agents. Fractions that contained purified TadA-GFPNb were concentrated, frozen in liquid N2, and stored at –80 °C.

#### TadA-λN

Protein was purified as described above for TadA-GFPNb.

#### pAG-TadA

Protein was purified as described above for TadA-GFPNb. Fractions with recombinant protein from the 1 mL heparin column were collected, concentrated, and further purified using a HiLoad 16/600 Superdex 200 pg SEC column (120 mL; Cytiva, cat # 28989335). Fractions that contained purified pAG-TadA were concentrated, frozen in liquid N2, and stored at –80 °C.

#### TadA

Protein was purified as described above for TadA-GFPNb.

The identity of purified proteins was confirmed using standard mass spectrometry analyses in the Proteomics and Metabolomics Shared Resource at Fred Hutch.

### Antibody

GFP antibody (HtzGFP-19F7) was acquired from the Memorial Sloan Kettering Cancer Center.

#### *In vitro* transcription

Synthetic DNA was purchased from IDT that encoded a T7 promoter (TAATACGACTCACTATAGG), the human β-globin 5′ UTR, a nanoLuciferase ORF, and the human β-globin 3′ UTR, with the 15-nucleotide boxB motif (GCC-CTGAAGAAGGGC) either inserted or not 56 nucleotides downstream of the nanoLuciferase stop codon. This was amplified with Phusion High-Fidelity DNA polymerase (ThermoFisher, cat # F530S) and primers CPL_184 and CPL_262. The PCR product was purified using a PureLink Quick PCR Purification Kit (Invitrogen, cat # K310002).

After amplification, the RNAs were *in vitro* transcribed using a MEGAscript T7 Transcription Kit (Thermofisher, cat # AM1334) for 3 hours at 37 °C and treated with Turbo DNase for 15 minutes at 37 °C. Transcribed RNA was purified via a GeneJET RNA Purification kit (ThermoScientific, cat # K7032), followed by Micro Bio-Spin P-6 Gel Columns (Bio-Rad, cat # 7326221) equilibrated 3x with 200 uL of water. The purity of RNA was determined by agarose gel electrophoresis and quantified by Nanodrop.

### Low throughput RNA editing assay

The reporter mRNAs (125 nM each, 250 nM total, containing either boxB or not) were refolded in water by heating to 95 °C for 2 minutes and then cooled slowly to RT. The refolded mRNAs were added to a reaction mixture containing TadA buffer (50 mM Tris-HCl pH 7.5, 25 mM KCl, 2 mM MgCl2). After incubating at 37 °C for 5 minutes, the indicated TadA recombinant protein (250 nM) was added. Reactions were incubated at 37 °C for 2 hours. RNA was purified via a GeneJET RNA purification kit and quantified by Nanodrop. Purified RNA (375 ng) and DNA primer CPL_372 (1 µM) were incubated together at 65 °C for 5 minutes and annealed on ice. RNA was reverse transcribed using Maxima Reverse Transcriptase (ThermoFisher, cat # EP0742) at 50 °C for 30 minutes followed by 85 °C for 5 minutes. The resulting cDNA was amplified by PCR as described above using primers CPL_373 and CPL_374. The PCR products were purified and analyzed by nanopore sequencing (Plasmidsaurus). FASTQ files containing basecalled nanopore reads were sorted into populations that either contained the boxB element or did not. In both populations, the 40 nucleotides immediately 3′ of the boxB motif insertion were compared against the non-edited reference sequence to identify adenosine to guanosine substitutions. Sequences with inserts or deletions were discarded.

### BoxB reporter oligo library design

Oligos for cloning the *in vitro* reporter pool were designed with a custom script to generate a comprehensive library of sequence variants. Each oligo was composed of a 5′ T7 promoter (TAATACGACTCACTATAGG), a forward handle (TGGCTTCGTTGTTGTGCT), a variable spacer region (TTTGTGTTCTCTTGTTCGTTCTG-GTTCGTT), a recorder region (TAGAATTACACCATAAT), and the 19-nucleotide boxB stem loop (GGGCCCT-GAAGAAGGGCCC), which extends the 15-nucleotide motif of the nanoLuciferase reporter by two G:C base pairs, with additional short buffer sequences flanking the variable regions. Ten-nucleotide barcode sequences devoid of A and separated by a Hamming distance of 2 were used to uniquely tag every oligo. To generate spacer variants, the full-length spacer sequence was truncated in two-nucleotide increments, creating a set of fragments; for each truncation, the spacer was split into 5′ and 3′ segments, and the fixed recorder region was inserted between these fragments to form a “spacer target” sequence. For randomizing the recorder region, all non-A nucleotides within the recorder were independently randomized in groups of 5 nucleotides to yield two distinct sets of target sequences. Each randomized spacer target was then incorporated into two oligo orientations by appending the boxB stem loop together with its buffer on either the 5′ or the 3′ side. The boxB stem loop was randomized in six windows of 3 or 4 nt, covering every single substitution and combinations within those windows. The final oligo sequences are assembled by concatenating the T7 promoter, the forward handle, the designed variable region (incorporating spacer, recorder region, and boxB stem loops with their buffers), the barcode buffer with the assigned barcode, and a reverse transcription handle (GCTGGCTTCTGTTCCGTTTG). This oligo pool of 96 designs, each 136 nucleotides long and together specifying 69,888 distinct sequences, was ordered from IDT as oPool oAS2176. See Supplementary Table 4 for the full list.

Oligos for the *in vivo* reporter pool with randomized boxB stem loops were designed as above and ordered from IDT as oPool oRB262, comprising 32 designs of 117 nucleotides that together specify 4,352 distinct sequences. See Supplementary Table 5 for full list.

Oligos for the λN site saturation mutagenesis library were designed by replacing one codon at a time with NNN at each of the 22 positions. The oligos were ordered from IDT as oPool oRB275, comprising 22 designs of 112 nucleotides that together specify 1,408 position-codon combinations. See Supplementary Table 6 for full list.

### High throughput *in vitro* RNA editing assay

CPL_303 (10 µM), the T7 promoter top strand, was pre-annealed to the single-stranded oligo pool oAS2176 (10 µM) to reconstitute a double-stranded promoter, in annealing buffer (20 mM Hepes pH 7.5, 100 mM KCl, and 2 mM MgCl2) by heating to 70 °C then cooling by 0.2 °C/s. The pool was transcribed and treated with DNase as described above. RNA was purified by phenol:chloroform extraction and ethanol precipitation.

The RNA libraries (250 nM total concentration) were added to TadA buffer along with any non-TadA protein, namely λN-EGFP or the anti-GFP antibody, at 125, 250, or 500 nM. After incubating at 37 °C for 5 minutes, the TadA enzyme or TadA fusion was added at the same concentration. Component-omission controls lacked the antibody, λN-EGFP, or TadA, with the remaining components held at 250 nM. If the reaction contained the anti-GFP antibody, it was instead incubated for 1 hour at room temperature before TadA recombinant protein addition as previously described (Xiao et al. 2024a). The reactions were incubated at 37 °C for 0.5, 1, or 2 hours; split-recruitment reactions using pAG-TadA were additionally incubated at 25 °C or 13 °C for 2 hours. RNA was purified by phenol:chloroform extraction and ethanol precipitation.

### Cell culture

HEK293T cells were cultured in Dulbecco′s modified Eagle medium (DMEM 1X, with 4.5 g/L D-glucose, + L-glutamine, -sodium pyruvate, Gibco 11965-092) supplemented with 10% FBS (Thermo 26140079) and passaged using 0.25% trypsin in EDTA (Gibco 25200-056). Cells were grown at 37 °C in 5% CO2. Cell lines were confirmed to be free of mycoplasma contamination.

### Integration of plasmid libraries into landing pad cell lines

hsAS126.3 (HEK293T *attP* Cas9*) cells (Nugent et al. 2024) were seeded to 80% confluency in one 10 cm dish per library. 9.6 μg of *attB\**-containing reporter library plasmid (pAS476, pAS507, or pAS517) and 2.4 μg of Bxb1 expression vector (pAS344) were transfected per 10 cm dish using FuGENE HD reagent (Promega), with 36 μl of reagent in 600 μl of Opti-MEM per dish. Each library was transfected into a single 10 cm dish then expanded into 15 cm dishes 48 hours post-transfection. Cells were selected with 2 μg/ml puromycin, added 72 hours post-transfection. Puromycin selection was ended after 8 days, or 6 days for the pAS517 pool, and cell pools were contracted back into a 10 cm dish. After puromycin selection, medium was replaced with medium containing 2 μg/ml doxycycline to induce the reporter and, where present, the editor. The resulting pools were hsAS206 (pAS476), hsAS208 (pAS507), and hsAS209 (pAS517). The hsAS208 no-editor pool received the same 72-hour doxycycline treatment as the editor-expressing pools. Cellular boxB editing was measured in hsAS206 and hsAS208, and the λN mutational scan in hsAS209.

### Library mRNA extraction

Library mRNA was harvested after 72 hr of doxycycline induction of TadA and boxB reporters from one 50-75% confluent 10 cm dish. Each 10 cm dish was treated with 1 ml 0.25% trypsin, and neutralized with 5 ml DMEM media. Cells were pelleted from 1/3 of this cell suspension and resuspended in 1 ml Trizol reagent (Thermo). Total RNA from these lysates was then extracted using the Direct-zol RNA Miniprep kit (Zymo) following the manufacturer’s protocol.

### High throughput sequencing of boxB reporters

2.3-7 μg of total RNA from *in vivo* libraries or 25-200 ng RNA from each *in vitro* enzymatic reaction was reverse transcribed into cDNA in a 50 μl reaction using Maxima H Minus reverse transcriptase (Thermo) and RT primer oRB213, which also contains a 7 nt UMI, at 25 °C for 10 minutes, 55 °C for 60 minutes, and 85 °C for 10 minutes. A 20-50 μl PCR was performed using Phusion Flash High-Fidelity PCR Master Mix (Thermo) for 6-24 cycles of 98, 65, and 72 °C for 20 seconds each, with cDNA template comprising 1/20th of the final volume, and with oPN776 as the reverse primer. The cycle number for each library was chosen from a SYBR Green qPCR of the same reaction. Indexed forward primers were used to enable pooled sequencing of all samples (one of oRB218-oRB225 or oRB287-oRB302). All of these PCR reactions generated a 192 bp amplicon that was cut out from a 2% agarose gel and cleaned using the Zymoclean Gel DNA Recovery Kit (Zymo). The boxB reporter libraries were sequenced on an Illumina NextSeq 2000 using custom sequencing primers oRB214 for Read 1 (79 nt), oRB215 for Read 2 (7 nt), and oRB217 for indexing (7 nt).

### High throughput sequencing of λN variant libraries

Total RNA from the hsAS209 pool was reverse transcribed as above using RT primer oRB308, which binds the T7 region of the reporter mRNA and carries a 7 nt UMI. The cDNA was amplified with oRB308 and the indexed forward primer oRB302 using Phusion Flash High-Fidelity PCR Master Mix for 24 cycles of 98, 65, and 72 °C for 20 seconds each, giving a 240 bp amplicon. To link each 20-nt barcode to its λN insert, a 298 bp amplicon was generated from the pAS499 plasmid pool with oRB309 and oRB308 for 12 cycles of 98, 63, and 72 °C for 20 seconds each. Both amplicons were cut out from an agarose gel, cleaned, quantified by Qubit, and sequenced single-end on an Illumina MiSeq with a 6 nt sample index.

### Computational analyses

Analyses were run as Snakemake workflows (Köster and Rahmann 2012) in Singularity containers, with samples, coordinates, and analysis selections specified in versioned CSV annotations. The analysis repository is https://github.com/rasilab/tada_boxb. Workflows used ghcr.io/rasilab/r_python:2.4.2 (Python 3.12.8, NumPy 1.26.4, pandas 2.3.1, SciPy 1.16.0, Matplotlib 3.10.3, Biopython 1.85), except the in vitro read counter, which used ghcr.io/rasilab/python:1.0.0 (Python 3.9.15), and the published-data reanalysis, which used ghcr.io/rasilab/hisat2_3n:1.0.1. All containers used in these workflows are publicly available at https://github.com/orgs/rasilab/packages.

### boxB reporter counting

A 10-nucleotide reporter barcode identified the mutagenesis window, recorder background, and recorder placement of each read. Variable and recorder bases were extracted at the coordinates specified for each barcode. Recorder positions were reported in the RNA orientation; an A-to-I edit appears as T-to-C on the sequenced complementary strand. Reads were assigned by exact barcode match, and their invariant flanking sequences were checked. Reads containing a reverse-primer handle before its designed position were rejected as truncated amplicons. Reads with an ambiguous base in the UMI or variable insert were discarded. Molecules were deduplicated on the seven-nucleotide UMI within each sample, reporter barcode, and insert. For cellular libraries, the reporter and UMI reads were processed together, with matching read identifiers and valid FASTQ records required. Recorder bases below Phred 30 were masked as N. Each site’s editing percentage was calculated as 100 times its edited molecule count divided by the number of molecules callable at that site. Stem-disrupting controls comprised the four annotated substitutions in each of windows 4–6 and 14–16. In vitro summaries used wild-type controls from all six windows and excluded the zero-spacer 5′ libraries for windows 1–3 and 7–10 because of read-quality defects, leaving 22 distinct wild-type constructs; cellular summaries retained all 24. The six mutagenesis windows contain 760 distinct full-length boxB sequences including wild type. Sequence effects were calculated by pooling edited and callable counts across retained placements and windows representing the same full sequence. Editing at individual boxB adenosines was calculated after excluding mutagenesis windows overlapping the measured adenosines.

### λN barcode linkage and variant counting

The λN library contains 22 positions × 64 codons, corresponding to 1,387 distinct inserts after identical wild-type designs were collapsed. Barcode and insert identities required Phred 30 at every identity base and a match to a complete oriented insert sequence. Constant-region matching permitted C at an editable reference T. Linkage required at least three reads supporting designed inserts and at least 90% designed-insert purity per barcode; barcodes with a one-substitution neighbour were excluded. Figure analyses required a minimum of five designed linkage reads before RNA counts were pooled. Each site’s editing percentage was calculated as 100 times the summed edited reads divided by the summed reads callable at that site.

Codon-level analyses retained distinct inserts with the exact wild type counted once. Amino-acid analyses pooled codons and barcodes for each substitution, pooled synonymous variants into one wild-type reference, and grouped stop codons by position. Position profiles averaged covered missense rates equally, while stop profiles pooled counts within each position.

### Reanalysis of published rABE sequence contexts

Fifteen RNA-sequencing samples from Lin et al. (Lin et al. 2023) covering PUM1–rABE, PUM2–rABE, unfused rABE, rbFOX2–rABE, and an uninduced control were aligned with HISAT-3N in A-to-G conversion mode against the unmasked GRCh38 primary assembly with Ensembl 108 splice annotations, using forward RNA-strand specification and repeat mode limited to 1,000. Sites were retained if they had at least one converted read and more than 10 total reads. Contexts were oriented to the edited strand, and N-containing contexts were excluded. Mean site-level editing fractions were calculated for each context by pooling retained sites across three replicates.

### Statistical comparisons

For in vitro and cellular WT–MUT comparisons, the four mutants sharing a wild-type barcode were averaged first, then compared with that wild type using paired two-sided t-tests across eight matched window and recorder-placement groups. Each wild-type control contributed once to each test. Bars summarize all retained constructs; tests use the matched stem-window groups. For individual boxB adenosines (Supplementary Figure 1B), the plotted 15 WT and 32 MUT constructs were instead compared directly using unpaired two-sided Welch t-tests. λN codon-class distributions were compared using two-sided Mann–Whitney U tests on variant-level mean editing percentages. Amino-acid substitution rates were compared with pooled synonymous wild type using two-sided Student t statistics, with variances inflated by barcode-level dispersion estimated from wild type and degrees of freedom equal to the number of contributing barcodes minus one (minimum one). Position-mean tests combined the variance across covered missense substitutions with the wild-type reference variance, using the number of substitutions minus one degrees of freedom. The λN position and substitution screens were corrected separately by the Benjamini–Hochberg procedure within each recorder site, and significance required a BH-adjusted P value below 0.05 at both sites. Pearson correlations were calculated on the plotted values, using log₂ values for logarithmic axes. For Figure 1C, the 95% confidence intervals describe read-sampling uncertainty, calculated with NumPy’s default random generator using seed 20261004.

### Use of generative artificial intelligence

We used OpenAI GPT (GPT-6 Astra and GPT-5.6 Sol) and Anthropic Claude (Opus 5 and Sonnet 5.5) models for data analysis and editing of this manuscript. All content in the manuscript and in the associated GitHub repository was manually reviewed for correctness by the corresponding authors.

---

**Supplementary Figure 1.**
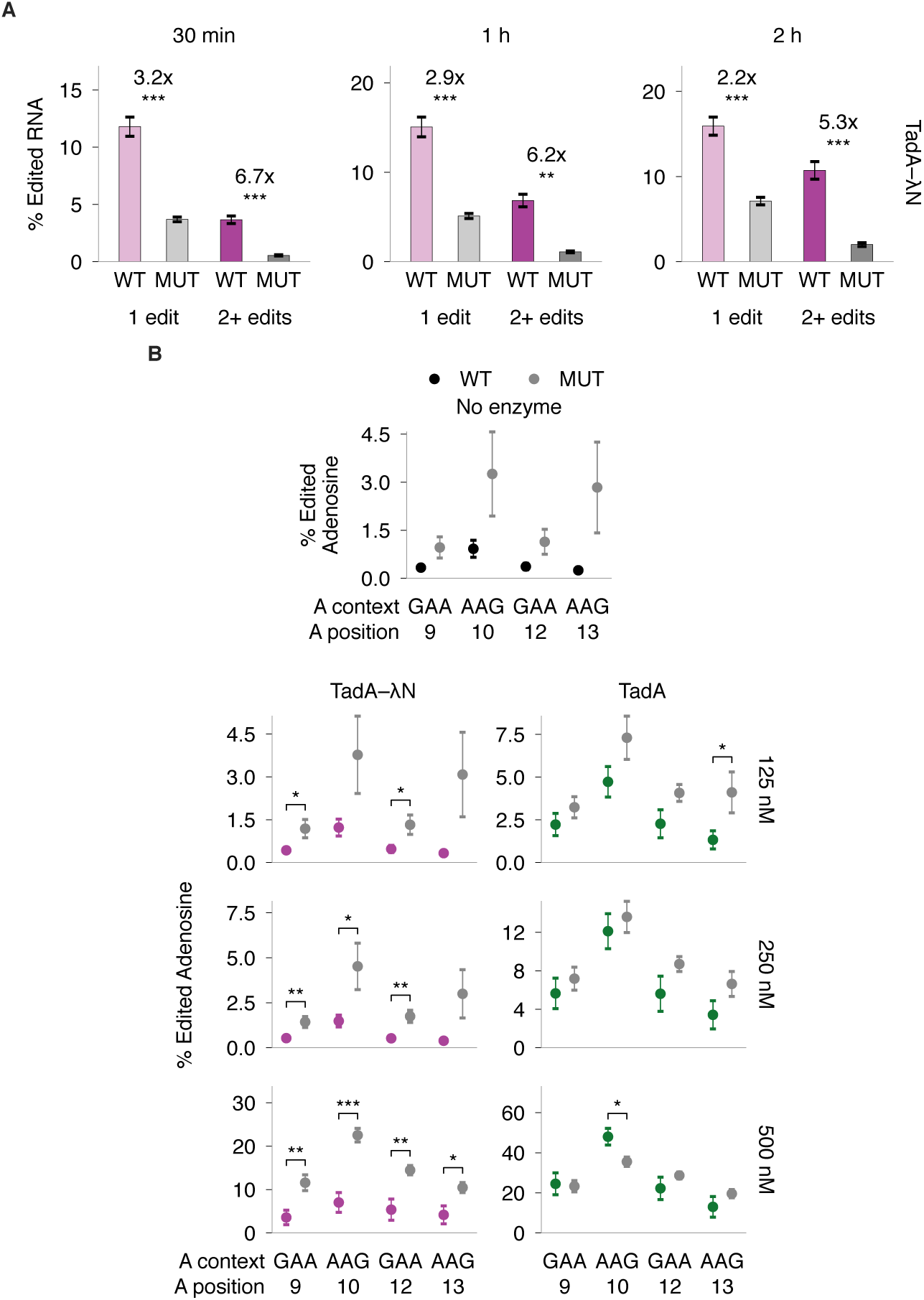
**(A)** Percentage with one or 2+ recorder edits after 30 min, 1 h, or 2 h with 250 nM TadA-λN. Bars show means ± SEM across 22 WT or 32 MUT constructs. Fold changes compare WT with MUT (paired two-sided t-tests). **(B)** Percentage of callable molecules edited at each of the four boxB adenosines, named below each plot by sequence context and position. The top plot is the no-enzyme control; the grid below shows TadA-λN and TadA at the indicated concentrations. Points show means ± SEM across 15 WT or 32 stem-disrupted (MUT) constructs. Brackets compare WT with MUT (unpaired two-sided Welch t-tests). All reactions used 250 nM RNA pool at 37 °C; B was incubated for 2 h. *: P < 0.05, **: P < 0.01, ***: P < 0.001; no star indicates P ≥ 0.05.

**Supplementary Figure 2.**
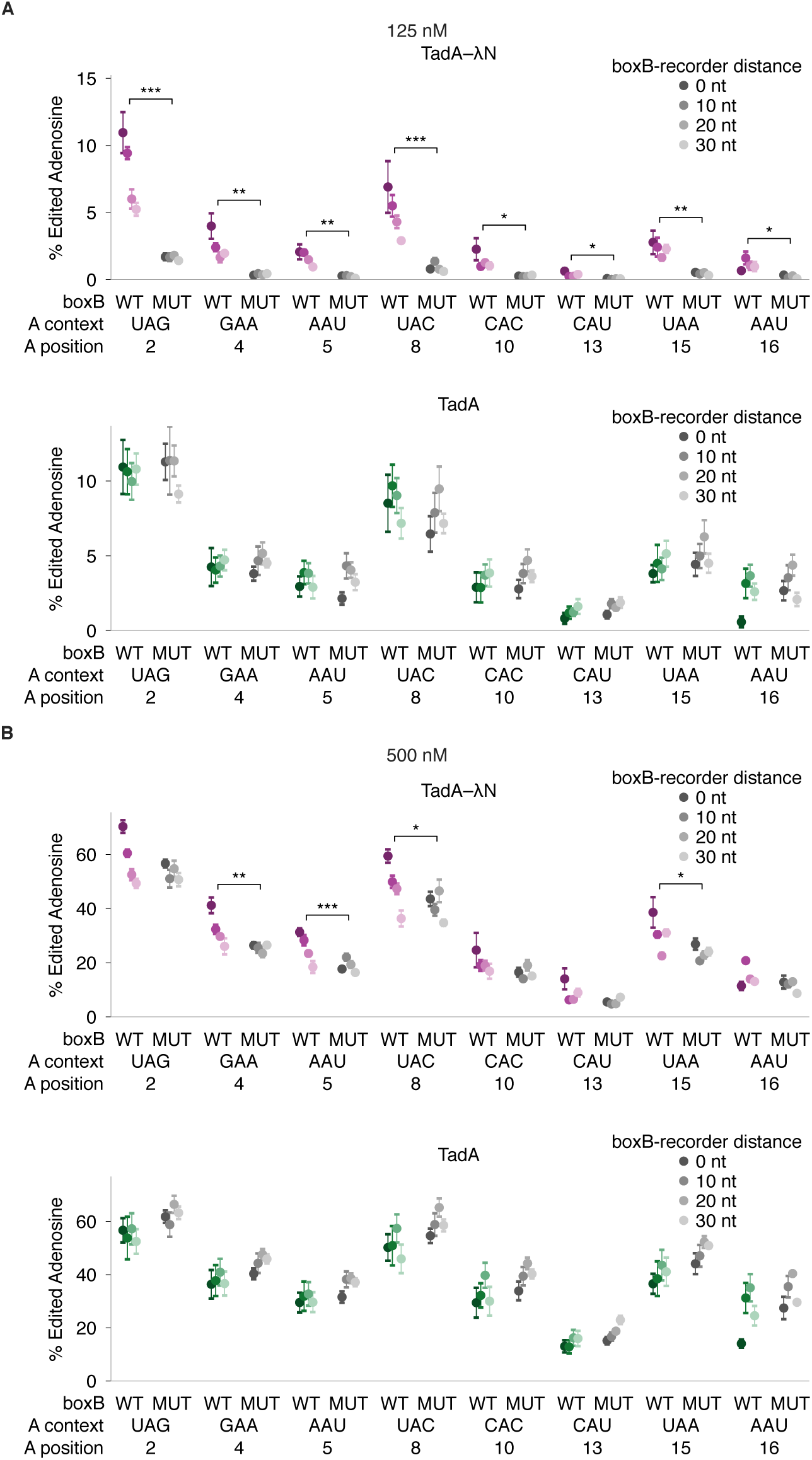
**(A,B)** Editing at each recorder adenosine versus boxB–recorder distance, at 125 nM **(A)** and 500 nM **(B)** enzyme. Each panel shows TadA-λN above TadA. Points show means ± SEM across four to six WT or eight MUT constructs per distance. All reactions used 250 nM RNA pool at 37 °C for 2 h. Brackets compare WT with MUT at each site; only significant comparisons are drawn. Tests use paired two-sided t-tests across eight matched window and placement groups. *: P < 0.05, **: P < 0.01, ***: P < 0.001; no star indicates P ≥ 0.05.

**Supplementary Figure 3.**
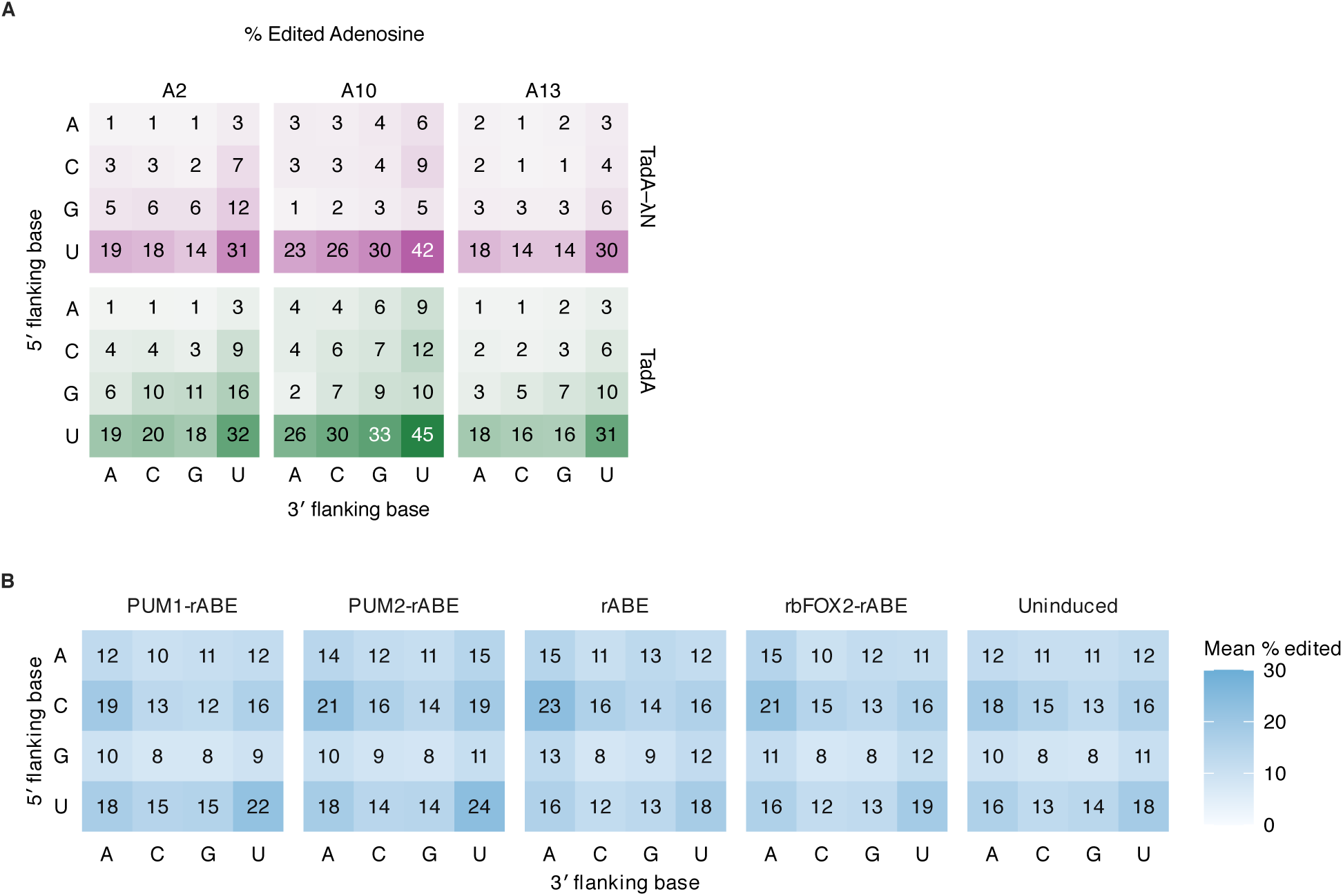
**(A)** Percentage of molecules edited at recorder positions A2, A10, and A13 by flanking context, for TadA-λN (top) and TadA (bottom). Each heat map pools all four boxB–recorder distances and 64 sequence backgrounds per distance, as in Figure 2E, and all six share one color scale. Color intensity varies in proportion to this percentage. Reactions used 250 nM enzyme and 250 nM RNA pool at 37 °C for 2 h. **(B)** Mean percentage of edited reads by flanking context in published rABE RNA sequencing (Lin et al. 2023), for PUM1-rABE, PUM2-rABE, unfused rABE, rbFOX2-rABE, and the uninduced control. Each value is the mean across edited sites that have at least one edited read and more than ten total reads, pooling three replicates per condition.

**Supplementary Figure 4.**
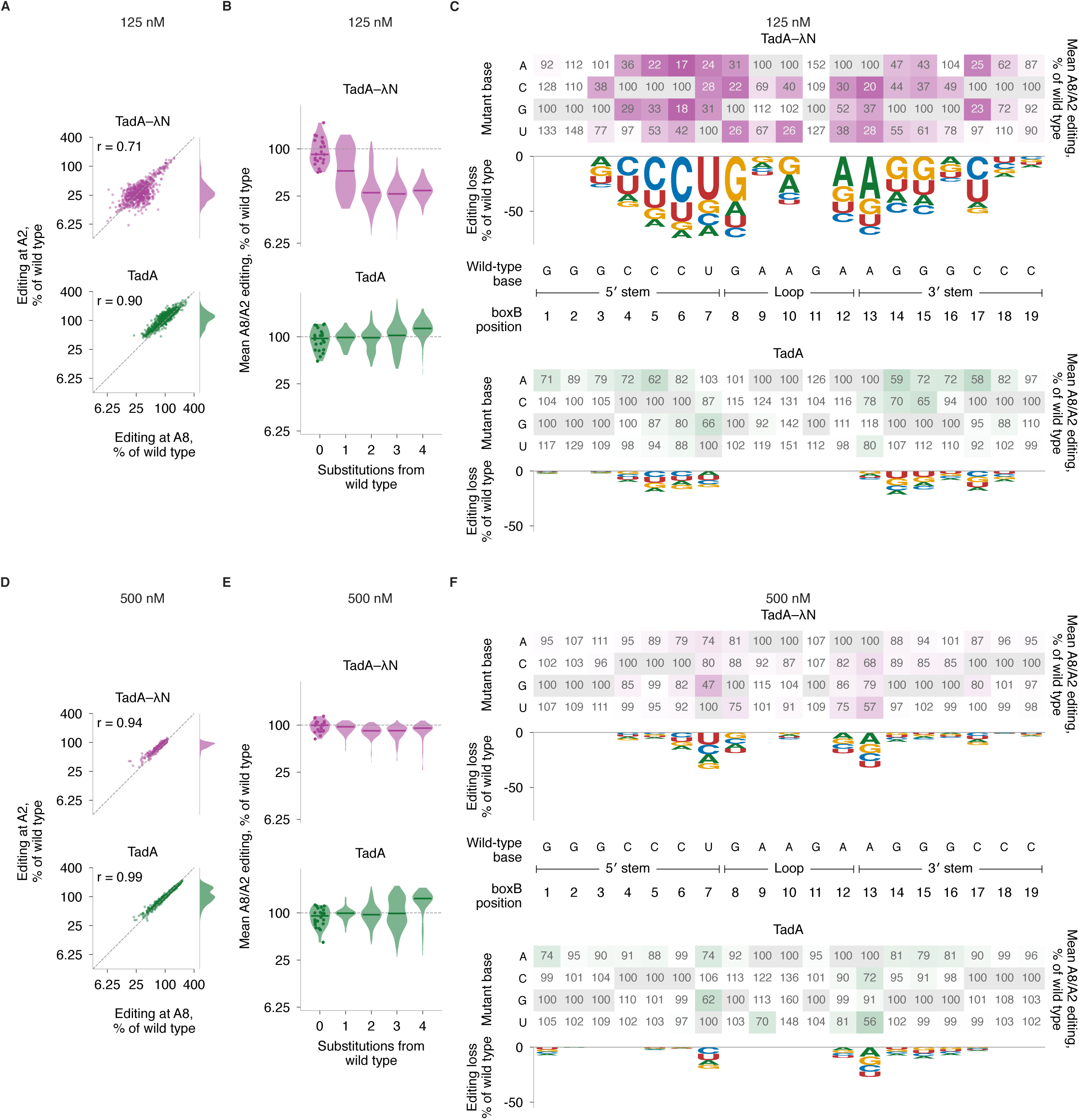
**(A)** Editing at recorder adenosines A2 and A8 for the 759 variants other than wild type, each normalized to its matched wild type, as in Figure 3B, at 125 nM. **(B)** Editing averaged across A8 and A2 as a function of the number of substitutions from wild-type boxB, as in Figure 3C. **(C)** Editing averaged across A8 and A2 of each single boxB mutant as a percentage of its matched wild type, as in Figure 3D. Gray cells are the wild-type base. Logos show the mean loss across the three substitutions at each position, with letters scaled by the editing each base retains; positions without a net loss have no stack. **(D-F)** The same three displays at 500 nM. All reactions used 250 nM RNA pool at 37 °C for 2 h.

**Supplementary Figure 5.**
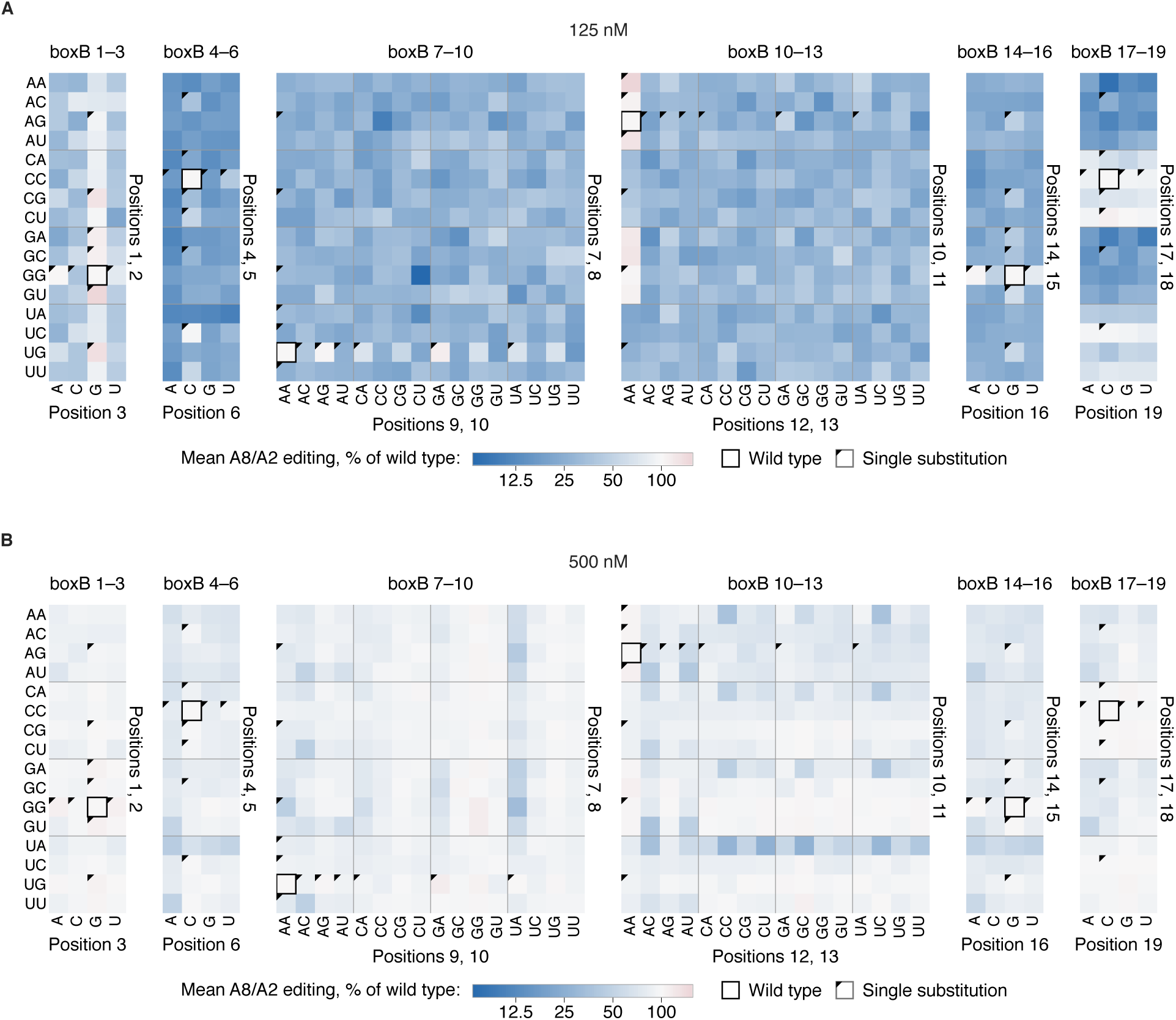
**(A,B)** Editing averaged across A8 and A2 for every variant of all six randomized windows, as in Figure 3F, at 125 nM **(A)** and 500 nM **(B)**. All reactions used 250 nM RNA pool at 37 °C for 2 h.

**Supplementary Figure 6.**
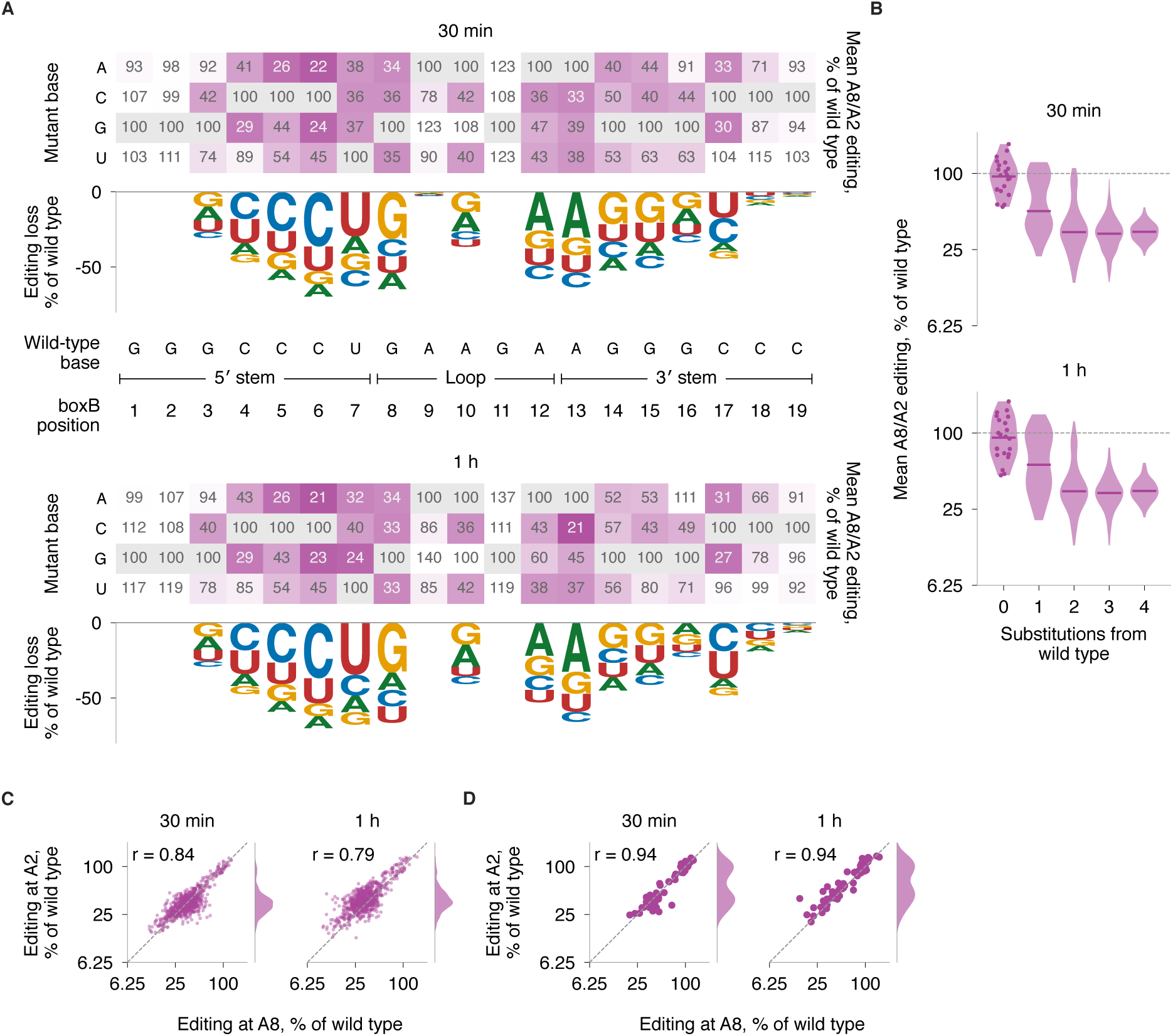
**(A)** Editing averaged across A8 and A2 of each single boxB mutant as a percentage of its matched wild type, as in Figure 3D, after 30 min and 1 h. **(B)** The same editing as a function of the number of substitutions from wild-type boxB, as in Figure 3C. **(C)** Editing at recorder adenosines A2 and A8 for the 759 variants other than wild type, as in Figure 3B. **(D)** The same display for the 57 single boxB mutants. All reactions used 250 nM TadA-λN and 250 nM RNA pool at 37 °C.

**Supplementary Figure 7.**
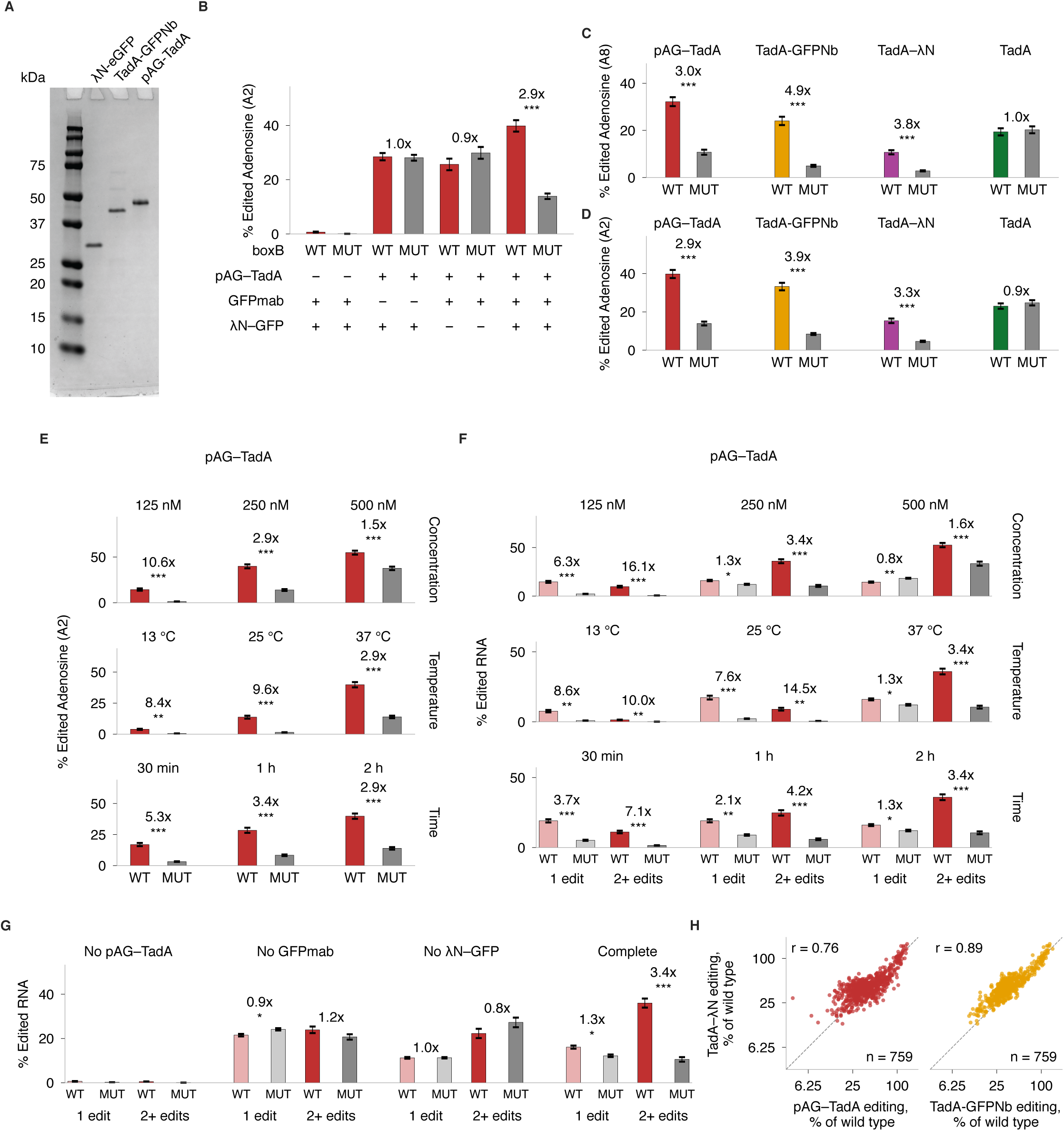
**(A)** SDS-PAGE analysis of purified λN-EGFP, TadA-GFPNb, and pAG-TadA, visualized by Coomassie staining. **(B)** A2 editing with the indicated components omitted, as in Figure 4C. **(C,D)** Editing at recorder adenosine A8 **(C)** and A2 **(D)** of WT or stem-disrupted (MUT) boxB by pAG-TadA, TadA-GFPNb, TadA-λN, and TadA. **(E)** A2 editing by pAG-TadA as a function of concentration, temperature, and incubation time, as in Figure 4D. Concentration applies to all recruitment components together. **(F)** Percentage of unique RNA molecules with one or 2+ recorder edits for WT or stem-disrupted (MUT) boxB, under the pAG-TadA conditions in E. **(G)** Percentage of unique RNA molecules with one or 2+ recorder edits with the indicated components omitted, as in Figure 4C. Bars in B–G show means ± SEM across 22 WT or 32 MUT constructs. Fold changes compare WT with MUT (paired two-sided t-tests). B, C, D, and G used 250 nM of each component at 37 °C for 2 h; E and F vary one condition at a time from that reaction. **(H)** Matched-variant editing effects for direct TadA-λN versus pAG-TadA or TadA-GFPNb recruitment, with every component at 250 nM, 37 °C for 2 h. Values are normalized to matched wild type and averaged across A8 and A2 on a log_2_ scale. The dashed line marks equal editing; r is the Pearson correlation of log_2_ ratios. Each panel contains all 759 non-wild-type variants. *: P < 0.05, **: P < 0.01, ***: P < 0.001; no star indicates P ≥ 0.05.

**Supplementary Figure 8.**
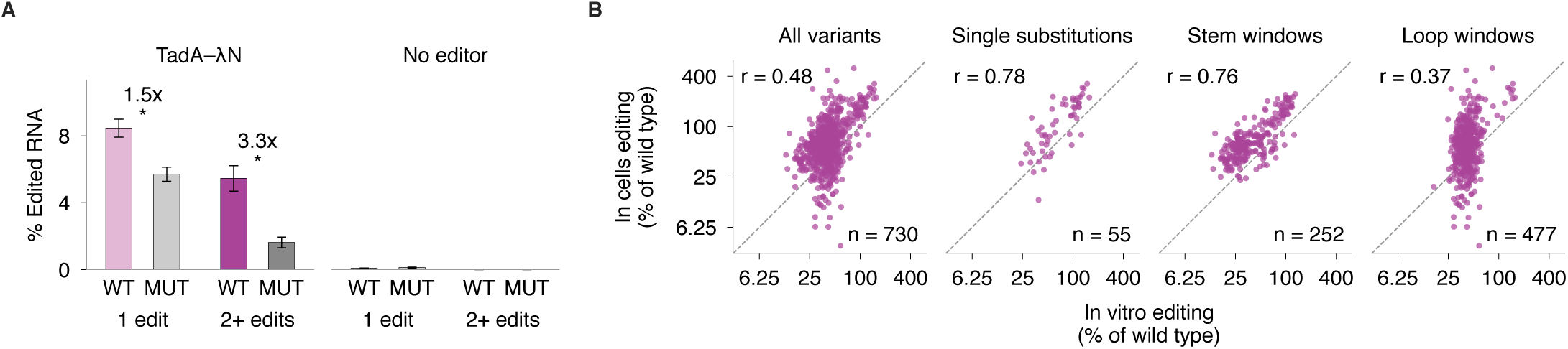
**(A)** Recorder editing with TadA-λN or no editor, as in Figure 5C. Bars show the percentage of unique RNA molecules with one or 2+ (2–8) edits for WT or stem-disrupted (MUT) boxB, as means ± SEM across 24 WT or 32 MUT constructs. Fold changes compare WT with MUT (paired two-sided t-tests). *: P < 0.05. **(B)** Editing in cells versus the 250 nM TadA-λN reaction in vitro, normalized to matched wild type and averaged across A8 and A2 on a log_2_ scale. The four plots show all variants, single substitutions, stem-window variants, and loop-window variants. The dashed line marks equal editing; r is the Pearson correlation of log_2_ ratios, and n is the number of variants plotted. Thirty of the 760 designed sequences were omitted because cellular editing was zero at one or both recorder sites.

